# Rapture-ready darters: choice of reference genome and genotyping method (whole-genome or sequence capture) influence population genomic inference in *Etheostoma*

**DOI:** 10.1101/2020.05.21.108274

**Authors:** Brendan N. Reid, Rachel L. Moran, Christopher J. Kopack, Sarah W. Fitzpatrick

## Abstract

Researchers studying non-model organisms have an increasing number of methods available for generating genomic data. However, the applicability of different methods across species, as well as the effect of reference genome choice on population genomic inference, are still difficult to predict in many cases. We evaluated the impact of data type (whole-genome vs. reduced representation) and reference genome choice on data quality and on population genomic and phylogenomic inference across several species of darters (subfamily Etheostomatinae), a highly diverse radiation of freshwater fish. We generated a high-quality reference genome and developed a hybrid RADseq/sequence capture (Rapture) protocol for the Arkansas darter (*Etheostoma cragini*). Rapture data from 1900 individuals spanning four darter species showed recovery of most loci across darter species at high depth and consistent estimates of heterozygosity regardless of reference genome choice. Loci with baits spanning both sides of the restriction enzyme cut site performed especially well across species. For low-coverage whole-genome data, choice of reference genome affected read depth and inferred heterozygosity. For similar amounts of sequence data, Rapture performed better at identifying fine-scale genetic structure compared to whole-genome sequencing. Rapture loci also recovered an accurate phylogeny for the study species and demonstrated high phylogenetic informativeness across the evolutionary history of the genus *Etheostoma*. Low cost and high cross-species effectiveness regardless of reference genome suggest that Rapture and similar sequence capture methods may be worthwhile choices for studies of diverse species radiations.

## Introduction

The advent of high-throughput sequencing (HTS) technology has enabled biologists to generate genome-scale molecular data from a variety of organisms, creating new opportunities for conservation genetics (Shafer et al. 2015), phylogenetics (Lemmon and Lemmon 2013, McCormack et al. 2013), and molecular ecology (Ekblom & Gallindo 2011). As the capacity for HTS has increased, however, repositories of sequence data have become increasingly biased toward sequences from a minority of model organisms (David et al. 2019). Although non-model organisms represent fruitful study systems for answering basic questions in biology (Russell et al. 2017), deciding on appropriate methods for generating and handling genomic data for non-model species remains a challenge.

Whole-genome sequencing may still remain out of reach for large-scale studies of non-model organisms, and as such reduced-representation approaches have grown popular as effective means for answering many questions (da Fonseca et al. 2016, Meek and Larson 2019). Sequence capture or targeted sequence enrichment methods represent an attractive method for generating repeatable, high-coverage sequence data (Grover et al. 2012, Harvey et al. 2016). A hybrid method that uses restriction-associated DNA sequencing (RADseq) combined with targeted enrichment of a user-defined subset of hundreds to thousands of RAD loci, termed ‘Rapture’ (Ali et al. 2016) has great potential as a rapid and efficient method for generating repeatable high-throughput genomic data at low cost and high efficiency. Rapture assays have so far been developed and applied to salmon (Ali et al. 2016), Tasmanian devils (Margres et al. 2018), marine turtles (Komoroske et al. 2019), frogs (Peek et al. 2019), and sea lampreys (Sard et al. 2020). The application of Rapture has mainly focused on population genomics within species, although Rapture loci developed for one species have been shown to be useful for studying hybridization among closely related species (Peek et al. 2019) and across species within slowly-evolving lineages (Komoroske et al. 2019).

For both whole-genome and reduced-representation sequencing, high-quality reference genomes can be used to improve genotype calling accuracy, inference of demographic history, and identification of loci under selection (Manel et al. 2015, Brandies et al. 2019). For studies of non-model species, however, reference genomes may not be available for the particular species of interest. Assembling HTS data to heterospecific genomes of related species is a potential option when such genomes are available. However, simulation studies indicate that even small divergences (0.15% to 2%) between the heterospecific reference genomes and the conspecific genome of the species of interest can increase errors in polymorphism calling and in estimates of genetic diversity, particularly when read depths are low (Nevado et al. 2014). Still, the practice of assembling short reads to a reference genome from a closely related species is common, and other empirical studies have concluded that congeneric or confamilial reference genomes may be suitable for SNP discovery, at least in groups with highly conserved genomes (Galla et al. 2019).

Using a conspecific reference genome in every situation is ideal but likely infeasible, especially when studying highly diverse species radiations. Applying HTS to the study of diverse species radiations will be particularly useful for understanding the effects of environmental context on genome evolution and identifying links between genetic variation and adaptive traits. Indeed, whole genome sequencing as well as reduced representation sequencing of adaptive radiations has uncovered signatures of change in genome structure and selection in African cichlid fish (Brawand et al. 2014) and specific genetic loci associated with beak and body size variation in Darwin’s finches (Chaves et al. 2016). However, the cost of generating separate reference genomes for each species may be prohibitive, and making population genomic comparisons among species often necessitates assembling data to a single reference genome (as in Chaves et al. 2016). If using heterospecific reference genomes is unavoidable in studies of diverse species radiations, it is important to quantify the biases that using these genomes will create when working with different types of data.

Darters (subfamily Etheostomatinae) represent a species radiation with great potential for illuminating the biotic and abiotic mechanisms that generate biological diversity. Darters are one of the most diverse clades of freshwater fish in North America, consisting of approximately 250 currently described species that likely shared a common ancestor between 30 and 40 million years ago (Near et al. 2011). Darters exhibit sexually dimorphic coloration that varies substantially among species, and sexual isolation based on divergent sexual selection has likely contributed to diversification in this group (Mendelson 2003, Moran et al. 2017, Moran and Fuller 2018a, Moran and Fuller 2018b). Postzygotic barriers between many sympatric species are not complete and hybridization is common, leading to gene tree discord and detectable signatures of ancient and contemporary introgression (Bossu & Near 2013; Moran et al 2017; Moran et al. 2018). Darters are dispersal-limited and often restricted to small headwater streams, and as such allopatric diversification due to physical isolation also plays a large role in their diversification (Near and Benard 2004, Hollingsworth and Near 2009). In addition to driving diversification, physical isolation and micro-endemicity, as well as habitat degradation, have created conservation issues for many darter species, and a substantial proportion of darter species diversity is currently considered threatened or endangered (Jelks et al. 2008).

HTS has great potential for providing insight into the forces controlling diversification in darters as well as for landscape and conservation genomics. Darter research to date has been characterized by a patchwork of molecular methods, making the comparison and integration of data from different studies difficult. Most previous phylogenetic work in darters has focused on Sanger sequencing of a small number of mitochondrial and nuclear genes (Near et al. 2011), while conservation genetics, landscape genetics, and molecular ecology studies have mainly used microsatellite markers developed for single species but with some applicability across the clade (Tonnis 2006, Khudamrongsawat et al. 2007, Switzer et al. 2008, Gabel et al. 2008, Hudman et al. 2008, Saarinen and Austin 2010). Recent work has begun to incorporate HTS methods, employing single-digest RADseq (Moran et al. 2018, MacGuigan et al. 2019, Moran et al. 2020) and double-digest RADseq (ddRAD, Moran et al. 2017, George 2018) to investigate phylogeny, phylogeography, and reproductive barriers among species. While ddRAD and RADseq represent a huge leap forward in terms of the amount of data generated, these methods often increase the number of loci genotyped at the expense of missing data and low coverage (MacGuigan et al. 2019). As such, there is currently no published method for reproducibly generating data for a single consistent set of loci distributed across the genome for darters. Furthermore, a reference genome assembly has only recently become available for a single darter species (the orangethroat darter *Etheostoma spectabile*; Moran et al. 2020).

Here, we describe an efficient and inexpensive Rapture-based method for reliably and repeatably genotyping thousands of loci in darters. This method is based on a capture bait set developed from RADseq data for Arkansas darters (*Etheostoma cragini*), a species of conservation concern found in the Arkansas River and nearby drainages within the Great Plains. Previous work in this species has used microsatellite markers to examine factors influencing population structure and genetic diversity in the western portion of their range (Fitzpatrick et al. 2014). The capture bait set targets over 2000 loci and includes both putatively neutrally-evolving loci as well as loci showing some evidence of selection across this species’ range. We assess two different tiling schemes for these baits, targeting either one or both flanking regions adjacent to a restriction cut site. We assess the performance of this capture bait set in a large set (*n* > 1600) of individual Arkansas darters as well as for individuals of three additional species in the genus *Etheostoma*. We assess the effects of aligning to either the heterospecific *E. spectabile* genome (which likely diverged from *E. cragini* approximately 29 million years ago; Kelly et al. 2015) or to a novel conspecific *E. cragini* genome, and we also compare estimates of genetic diversity and population structure from Rapture to estimates from low-coverage whole-genome sequencing (WGS) data for a subset of *E. cragini* individuals. We ask the following questions to gauge the performance and applicability of the method: 1) How often are loci sequenced using the Rapture baits recovered at high coverage (>20x), and how many reads per individual are needed to attain high coverage?; 2) How much diversity is present within the set of Rapture loci for both the target species and for other darter species?; 3) Can the Rapture loci identify distinct population units within *E. cragini*?; and 4) Do the Rapture loci recover known phylogenetic relationships among and within species? We also demonstrate how the choice of data type (Rapture vs. WGS) and reference genome (heterospecific vs conspecific) affects inference of population genetic parameters and population structure.

## Methods

### Sampling

Dipnetting and electrofishing were used by Kansas Department of Wildlife, Parks, and Tourism personnel to collect 2,374 *E. cragini* individuals at 216 sites throughout Kansas in 2015-2016. Fin clips were taken from adults (>28mm) and whole specimens were collected for juveniles (<28mm). Samples were stored in 100% ethanol, shipped to Michigan State University (MSU), then stored in a freezer (−20°C) prior to analysis. In addition to the Kansas samples, whole *E. cragini* specimens were collected from six sites in Arkansas by the Arkansas Fish and Game Commission. Tissue samples and isolated DNA from *E. cragini* individuals collected by the Colorado Department of Parks and Wildlife were also available from a previous study (Fitzpatrick et al. 2014). Sample information is provided in Supporting Table 1.

**Table 1.** Cost of Rapture and WGS methods used in this study.

| Method | # samples | Pilot dataset | Baits | Library preparation | Sequencing | Total cost | Cost/sample | Cost/sample after bait design |
| --- | --- | --- | --- | --- | --- | --- | --- | --- |
| BestRAD (pilot) | 52 | N/A | N/A | \$733.00 | \$2,661.00 | \$3,394.00 | \$65.27 | |
| Rapture | 1900 | \$3,394.00 | \$3,600.00 | \$5,190.00 | \$13,305.00 | \$25,489.00 | \$13.42 | \$9.73 |
| WGS | 24 | N/A | N/A | \$2,616.00 | \$3,991.50 | \$6,607.50 | \$275.31 | |

To examine the efficacy of Rapture across darter species, we also obtained genetic samples from three additional darter species: rainbow darters (*E. caeruleum*) collected for a separate population genetic study in southwestern Michigan (Oliveira et al. 2020), *E. spectabile* specimens collected in the Salt Fork of the Vermillion River, Illinois, and fantail darter (*E. flabellare*) specimens collected in Fox Creek, Illinois.

### DNA extractions

DNA from a pilot set of 52 *E. cragini* individuals sampled from seven sites across the species’ range was extracted using Qiagen DNeasy Blood & Tissue kits (Qiagen, Hilden, Germany). These extractions were done with a 60 ul elution in Qiagen EB buffer and quantified using a Qubit (Thermo Fisher Scientific, Waltham, MA, USA). For high-throughput extractions, we used a KingFisher Flex DNA extraction system (Thermo Fisher Scientific) to extract DNA from 20 sets of 90 samples (1800 samples total). We included an overnight digestion step in which tissues were lysed in a 96-well PCR plate at a constant temperature of 55° C on an Eppendorf Mastercycler thermal cycler (Eppendorf, Hamburg, Germany), and we included 10 uL Proteinase K solution, 10 uL enhancer solution, 100 uL Qiagen Buffer EB solution, and approximately 10 mg tissue in each digest. We then used the MagMax whole blood protocol for an input volume of 200 uL and a final elution volume of 60 uL. We quantified DNA yield from high-throughput extractions using a PicoGreen assay (Thermo Fisher Scientific, Waltham, MA), with the six wells left unused in each plate used for assay standards and a negative control. High-throughput extractions included an additional 1635 *E. cragini* samples from Kansas, 60 *E. cragini* samples from Arkansas, 20 *E. cragini* samples from Colorado, and all seven *E. spectabile* and eight *E. flabellare* samples (*E. caeruleum* samples were already extracted). DNA extracted for this study from *E. cragini* covered a total of 232 collection sites (*n* = 2-10 per site; Supporting Table 1). Nearly all of these samples yielded high-quality DNA and were included in the Rapture genotyping analyses described below.

### Pilot RADseq library preparation & Illumina sequencing

Using the pilot set of 52 *E. cragini* samples, two initial RADseq libraries (consisting of 24 samples and 28 samples respectively) were prepared and submitted to the MSU core genomics facility for sequencing. We used the ‘BestRad’ protocol following Ali *et al.* 2016. Briefly, genomic DNA (100 ng) from each sample was digested with a restriction enzyme (Sbfl-HF) and indexed with a biotinylated RAD adapter. Pooled DNA was sheared to 500 bp fragments using a Covaris sonicator (Covaris, Woburn, MA, USA). Shearing efficiency was evaluated with a fragment analyzer. Dynabeads M-280 streptavidin magnetic beads (Thermo Fisher Scientific, Waltham MA, USA) were used to physically isolate the RAD-tagged DNA fragments. The DNA was then eluted in TE buffer and used in NEBNext Ultra DNA Library Prep Kit for Illumina (New England Biosciences, Ipswich, MA, USA) with no modifications. The two libraries were each sequenced with paired-end 150 bp reads on an Illumina HiSeq 4000 in separate lanes.

### Bioinformatic pipeline for pilot dataset

As the BestRad protocol can result in sequences with barcodes on either the forward or reverse reads, we used a Python script (Flip2BeRad, https://github.com/tylerhether/Flip2BeRAD) to flip sequences with barcodes on the reverse read. We then filtered out potential PCR clones and demultiplexed sequences using the clonefilter and process_radtags commands in Stacks v. 2.4 (Catchen et al. 2013; Rochette et al. 2019). Using the demultiplexed forward reads, we identified loci containing single-nucleotide polymorphisms (SNPs) in ipyrad (Eaton and Overcast 2020). Reads were filtered using ipyrad’s default quality thresholds and mapped to an early draft version of the *E. spectabile* genome. We retained an initial set of candidate loci that were genotyped in ≥75% of the pilot set of 52 *E. cragini* individuals and that contained SNPs with a minor allele frequency (MAF) > 0.05. Additionally, we only retained loci with SNPs that were called in at least two *E. cragini* individuals, which imposed an additional floor on MAF and removed SNPs called in only one individual due to sequencing error. We created a FASTA file for all loci that passed these allele frequency filters and aligned these sequences to the draft *E. spectabile* genome using bwa v.0.7.17 (Li and Durbin 2009). As bait capture is optimally efficient as long as sequences are >95% similar to baits (Arbor Biosciences, personal communication), any sequences that exhibited <95% similarity or aligned to multiple locations on the *E. spectabile* draft genome were removed from the candidate set. Because the *E. spectabile* draft genome contained many small scaffolds, we also removed any loci that were located on scaffolds smaller than 10kb, as it would be difficult to determine whether these loci were adjacent to any other loci in the final chromosome-level genome assembly.

To identify population clustering within the pilot samples, we conducted an exploratory PCA using the r package adegenet (Jombart 2008, Jombart & Ahmed 2011) (Supporting Figure 1). This preliminary analysis indicated that pilot samples clustered into three distinct groups. To identify potential signatures of selection in this initial set of candidate loci, we used the program BayeScan (Foll and Gaggiotti 2008), which takes a Bayesian approach to identify outlier loci with higher or lower F_ST_ values than expected by chance given population structure. We conducted an initial analysis using all populations. After this analysis showed a high average F_ST_ and an overabundance of lower-than-expected F_ST_ outliers, we re-ran the analysis using only populations in the mainstem Arkansas river (Supporting Figure 2). We used a false discovery rate of 0.05 to identify outlier SNPs in both datasets.

### Bait design

From the candidate set of RAD loci, we identified three different categories of potential baits to be used as targets for Rapture: (1) short loci (n=3,176), in which ipyrad identified a locus containing at least one SNP that was located on one side of the restriction cut site only; (2) long loci (n=249), consisting of paired loci that both contained a SNP and were located on either side of the cut site; and (3) outlier loci (n=29) identified by Bayescan. Long loci were initially chosen to assess stretches of homozygosity or as potentially more phylogenetically informative blocks of sequence. We obtained BED coordinates for all target loci on the *E. spectabile* draft genome and provided these coordinates and the draft genome to Arbor Biosciences (Ann Arbor, MI, USA). Arbor Biosciences designed and produced a set of 4,966 80-bp baits to capture all long and outlier target RAD loci, as well as 1,841 of the short target RAD loci, for a total of 2,119 Rapture loci. While most previous Rapture study designs have used 120-bp baits (Ali et a. 2016, Komoroske et al. 2019, Peek et al. 2019), we used 80-bp baits tiled in an overlapping manner along the target loci to increase capture efficiency (as in Sard 2020). For short and outlier loci, two baits were tiled along each locus (starting at the restriction site), meaning that approximately 40 bps in the center of each locus were covered by two baits and the regions flanking this central region were only covered by one bait. For the long loci, five baits were tiled across both regions flanking the restriction site (Figure 1), meaning that a much longer region (approximately 160 bp) was covered by more than one bait.

**Figure 1.**
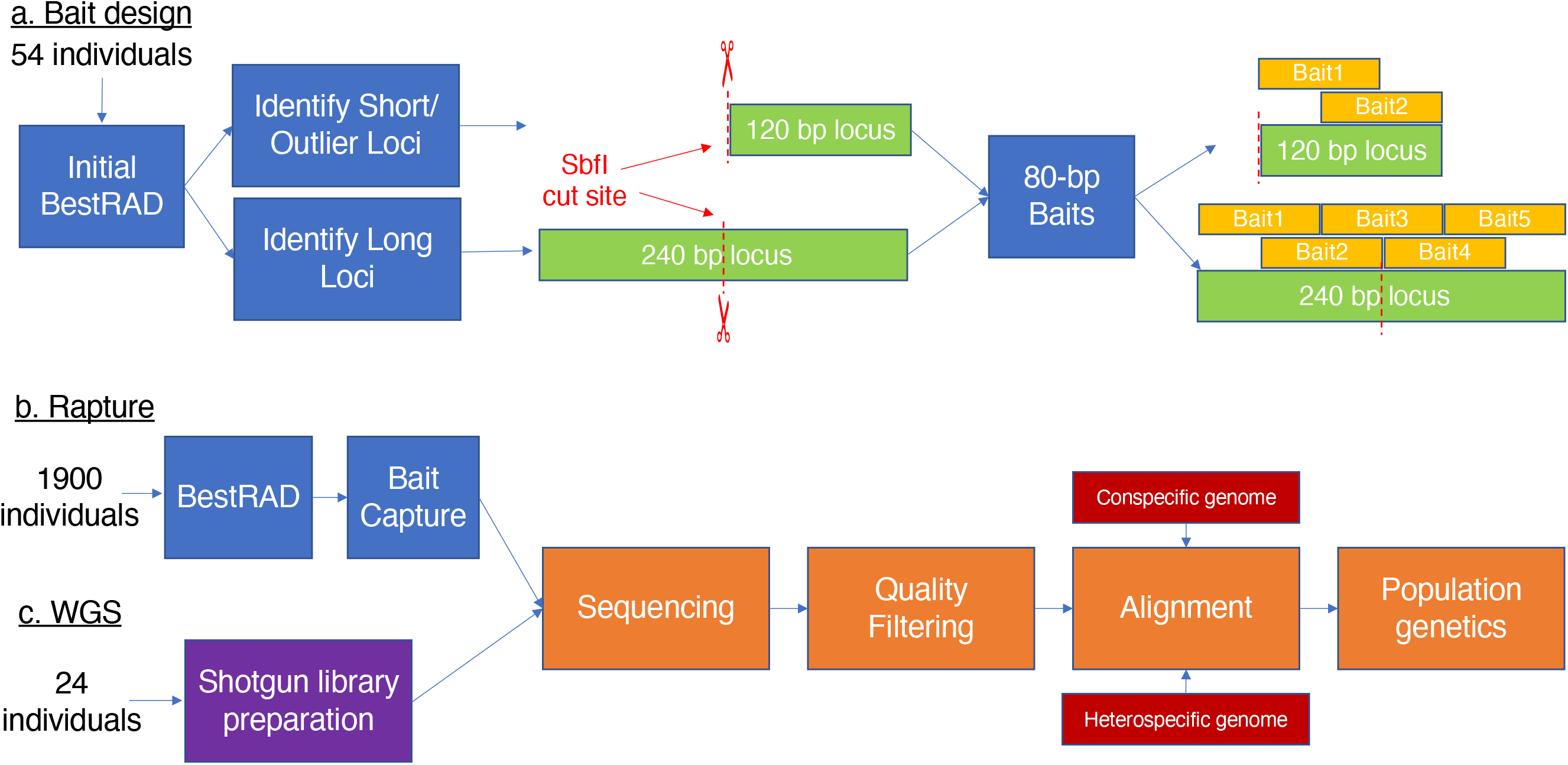
Flow chart showing procedures used to design *E. cragini* Rapture baits from BestRAD RADseq data, test the Rapture baits using a subsequent round of BestRAD RADseq, and compare the results of population genomic analyses using Rapture versus WGS data.

### E. cragini whole genomes

To compare population genetic statistics generated with Rapture to those generated using WGS, we produced a reference genome for *E. cragini* and conducted low-coverage whole-genome resequencing. We submitted *E. cragini* muscle tissue from two young-of-year fish of unknown sex raised at the John W. Mumma Native Aquatic Species Restoration Facility to Dovetail Genomics (Scotts Valley, CA, USA) to produce a high-quality reference genome for this species. Dovetail performed Illumina shotgun library preparation, paired-end 2×150 sequencing on an Illumina HiSeq X, and *de novo* assembly in Meraculous (Chapman et al. 2011) using a kmer size of 79. The assembly was refined using Chicago and Hi-C libraries, and scaffolds were constructed using HiRise (Putnam et al. 2016).

We submitted isolated DNA from 24 *E. cragini* samples for low-coverage whole-genome resequencing at the MSU Core Genomics center. Samples were chosen to include several individuals in each of several population clusters identified by Rapture (see below). We used the Illumina Coverage calculator (https://support.illumina.com/downloads/sequencing_coverage_calculator.html) to estimate the amount of sequencing needed to achieve ≥5x coverage based on genome size and an estimate of 20% duplicate sequences. These samples were submitted in two batches of 12, and each batch also contained four samples from another fish with a similar genome size (*Gambusia affinis*). As such, we used 75% of a lane of sequencing for each batch of 12 samples (1/16^th^ of a lane for each sample). Due to maintenance problems at MSU, the initial batch of sequencing produced fewer reads than expected. MSU sent the first batch of samples to the University of Michigan genomics core for additional sequencing and sent the second batch of samples to the Illumina FastTrack Sequencing Service Center for sequencing. All sequencing was performed on an Illumina HiSeq 4000.

We used FastQC (http://www.bioinformatics.babraham.ac.uk/projects/fastqc/) to assess sequencing quality across individuals. We used BWA v. 0.7.17-r1188 (Li and Durbin 2009) to align sequences to either the conspecific *E. cragini* genome or the heterospecific *E. spectabile* genome. We used samtools v.1.9 (Li et al. 2009) to filter out low-quality sequences and improperly paired reads, remove duplicates, and compute average coverage over the whole genome and over all covered sites for alignment to either the conspecific or heterospecific genomes.

### Rapture library preparation, sequencing, data processing pipeline, and quality control

We used the BestRAD protocol described above along with a sequence capture step that incorporated the Rapture bait sequences to conduct reduced-representation library preparation for 1900 individuals (1,855 *E. cragini*, 28 *E. caeruleum*, 8 *E. flabellare* and 9 *E. spectabile*). We aimed for a target DNA mass of 200 ng in 10 uL for the starting material in each reaction. For DNA samples with concentrations of 15-20ng/uL of DNA, we used 10uL total DNA. For samples with concentrations <15 ng/uL, we used a ThermoSavant DNA120 Speedvac (Thermo Fisher, Waltham, MA, USA) to dry down a sample volume containing 200ng and then resuspended in 10 uL 1x TE buffer. We performed library preparation in batches of four 96-well plates (containing 95 samples and one 1X TE blank), using the BestRAD barcode sequences and a plate-specific Illumina adapter for each plate. After BestRAD library preparation, we pooled all four plates and performed sequence capture using the protocol provided by Arbor Biosciences. Briefly, this involved performing a hybridization step at 65°C for at least 16 hours, isolating bait-target hybrids using streptavidin-coated magnetic beads and washing to remove non-target DNA, and performing PCR amplification of captured DNA for sequencing. We submitted these libraries for sequencing at the MSU Genomics Core facility in five batches of 380 samples each using paired-end 2×150 bp reads on an Illumina HiSeq 4000, using a single lane of sequencing for each batch. For each batch, we altered the number of cycles used for PCR amplification of libraries during the library preparation and the sequence capture steps in order to ensure a high enough concentration for sequence capture and sequencing, respectively. We used 12 cycles during library preparation for all libraries except libraries used in batch 3, where 11 cycles were used. For PCR amplification during sequence capture we used 12 cycles in the first 2 batches and 11 cycles in the three subsequent batches. We used the steps described above for BestRAD to process the raw data, and we used BWA to align reads to both the *E. spectabile* (v. UIUC_Espe_1.0, downloaded from NCBI; Moran et al. 2020) and the *E. cragini* reference genomes, and used samtools v. 1.7 (Li et al. 2009) to remove improperly paired reads. We generated two updated bed files by aligning the baits to each genome, merging all loci together into a single file, and creating a buffer (+500 bp from the 3’ end of the baits for short and outlier loci, +/− 500 bp on either side of the baits for long loci). We filtered all BAM files using these buffered regions before performing population genetic and phylogenetic analyses.

We evaluated data quality and potential differences caused by mapping to either a conspecific or heterospecific reference genome using multiple metrics, including the proportion of clonal reads per library preparation, the proportion of reads mapping to either reference genome per individual sample, and the proportion of mapped reads that overlapped the buffered Rapture loci per individual sample. We evaluated potential batch effects by comparing these metrics across Rapture batches. We also estimated individual-level coverage of Rapture loci using bedtools. We assessed two metrics of coverage: (1) the number of reads with any overlap for each buffered locus; and (2) per-base coverage of each buffered locus for a subset of individuals to examine how read depth changed with distance from the restriction site for different types of loci (long vs short).

### Population genomics, population structure, and selection

We used ANGSD v.0.928 (Korneliussen et al. 2014) to calculate genotype likelihoods for single-nucleotide polymorphisms (SNPs) based on aligned BAM files for Rapture and low-coverage WGS data. As we only had WGS data from *E. cragini*, we excluded the other species from this analysis. To create subsetted datasets with comparable numbers of total reads for comparing population genetic inferences between Rapture and WGS, we randomly selected 570 individuals (corresponding to the number of individuals sequenced on 1.5 lanes) from the Rapture dataset. We assumed bi-allelic SNPs when calculating genotype likelihoods and estimated major and minor allele frequencies for all SNPs. We used sites for which ANGSD detected a SNP with a *p*-value of < 1× 10^−6^, and we discarded SNPs with a minor allele frequency < 0.05 and SNPs which were genotyped in <50% of individuals (after Komoroske et al. 2018). We then used the program PCAngsd v.0.981 (Meisner and Albrechtsen 2018) to conduct downstream population genomic analyses. We first conducted principal component analyses (PCA) and calculated genotype probabilities on each of six datasets (WGS, full Rapture, and subsetted Rapture sequence sets, with each set aligned either the *E. spectabile* or *E. cragini* genome), with the optimum number of principal components determined by PCAngsd using a minimum average partial (MAP) test. We examined PCA results to evaluate evidence for batch effects in PCAngsd analyses. To obtain estimates of heterozygosity, we used PCAngsd to call genotypes using a probability threshold of 0.9. We compared matched individual heterozygosity values between data types (WGS or Rapture) and between data aligned to either the *E. spectabile* or the *E. cragini* reference genome. We also used PCAngsd to estimate individual admixture proportions for each individual and to perform a PCA-based scans for loci potentially under selection (i.e. loci exhibiting greater differentiation along PCs than expected by drift; Galinsky et al. 2016). We calculated *p*-values for the test statistics generated by PCAngsd selection scan using a one-tailed chi-squared test with one degree of freedom. To account for the large number of tests conducted for the selection scan, we set a conservative significance threshold for each dataset (Rapture and WGS) of 0.05 divided by the number of SNPs in the dataset. We compared within-species population structure and selection scan results along the first principal component axis between data types and reference genomes for all *E. cragini* individuals.

### Phylogenetics and phylogenetic informativeness

We compiled filtered BAM files aligned to the *E. cragini* genome from a subset 56 individuals (two individuals from *E. spectabile* and *E.flabellare*, two individuals from each of two sites for *E. caeruleum*, and two individuals from each of 19 sites covering the full distribution of *E. cragini*) and ran the ref_map.pl script in Stacks using the “populations: phylip_var_all” option and default parameter values to call SNPs and output PHYLIP-formatted concatenated multiple sequence alignments for each individual. We then used IQTREE (Nguyen et al. 2015) to construct a phylogenetic tree of all sequences. We used the default maximum likelihood model selection and tree search methods in IQTREE with 1000 bootstraps to calculate support values.

We converted this tree into a time-calibrated ultrametric tree using the R package ape (Paradis and Schliep 2019). We set estimated branching times for three splits based on a published study of darter evolution (Kelly et al. 2012) using the makeChronosCalib function to calibrate ranges of potential branching times for three interspecific splits. We set the root of the tree, identified here as the common ancestor of the clades *Oligocephalus*, *Psychromaster*, and *Catonotus*, to 24-34 million years ago. We also set the root of *Oligocephalus* (corresponding to the *E. caeruleum* – *E. spectabile* split in our tree) to 17.5-27.5 million years ago, and the common ancestor of *Psychromaster* and *Catonotus* (corresponding the *E. flabellare* – *E. cragini* split in our tree) to 16.5 – 26.5 million years ago. We then used the function chronos to construct a time-calibrated tree under three clock models (correlated, relaxed, and discrete). The correlated model had the highest likelihood and we used the tree calibrated using this model in all further analyses. We plotted the time-calibrated tree in ape and plotted the tips of the tree in space using the R package phytools (Revell 2012).

To calculate phylogenetic informativeness, we created separate PHYLIP files for each set of loci (short, long, and outlier) and concatenated all three into a single dataset and exported as a Nexus file using ape. We then input this alignment and the time-calibrated tree into the PhyDesign web interface (López-Giráldez and Townsend 2011). We examined the inferred net phylogenetic informativeness for each set of baits over the time period covered by the phylogeny (30 million years ago – present).

## Results

### E. cragini whole genomes

The *E. cragini* genome was similar in terms of contiguity and completeness compared to other published percid reference genomes, although it was smaller (643 Mb vs. greater than 850 Mb in all other percid genomes), contained less repetitive content, and exhibited a number of chromosomal rearrangements, especially relative to *E. spectabile* (Supporting Information 1, Supporting Figure 3). Coverage for resequenced *E. cragini* individuals varied based on the number of reads generated and the reference genome used. Shotgun sequencing for low-coverage WGS generated between 20.5 – 37.8 million read pairs per individual. Between 8.1%-15.5% of sequences were duplicates. Average read depth for covered sites (i.e. all sites with at least 1x coverage) and for all sites in each genome increased with the number of reads (Figure 2; *r* > 0.99 in all cases). Average read depth and depth of covered sites were highest when reads were aligned to the *E. cragini* genome and were almost identical, indicating that nearly all sites in the *E. cragini* genome assembly were covered at least 1x. Read depth progressively decreased by approximately 20% for covered sites and by approximately 44% for all sites when reads were aligned to the *E. spectabile* genome (Figure 2). While some of the decline in coverage over all sites may be attributable to the 30% greater length of the *E. spectabile* assembly (which is suggestive of a reduction in genome size for *E. cragini* relative to *E. spectabile*), lower coverage at sites with at least 1x coverage (presumably present in both genomes) also suggests loss of sequencing information resulting from poor alignment to the heterospecific reference genome.

**Figure 2.**
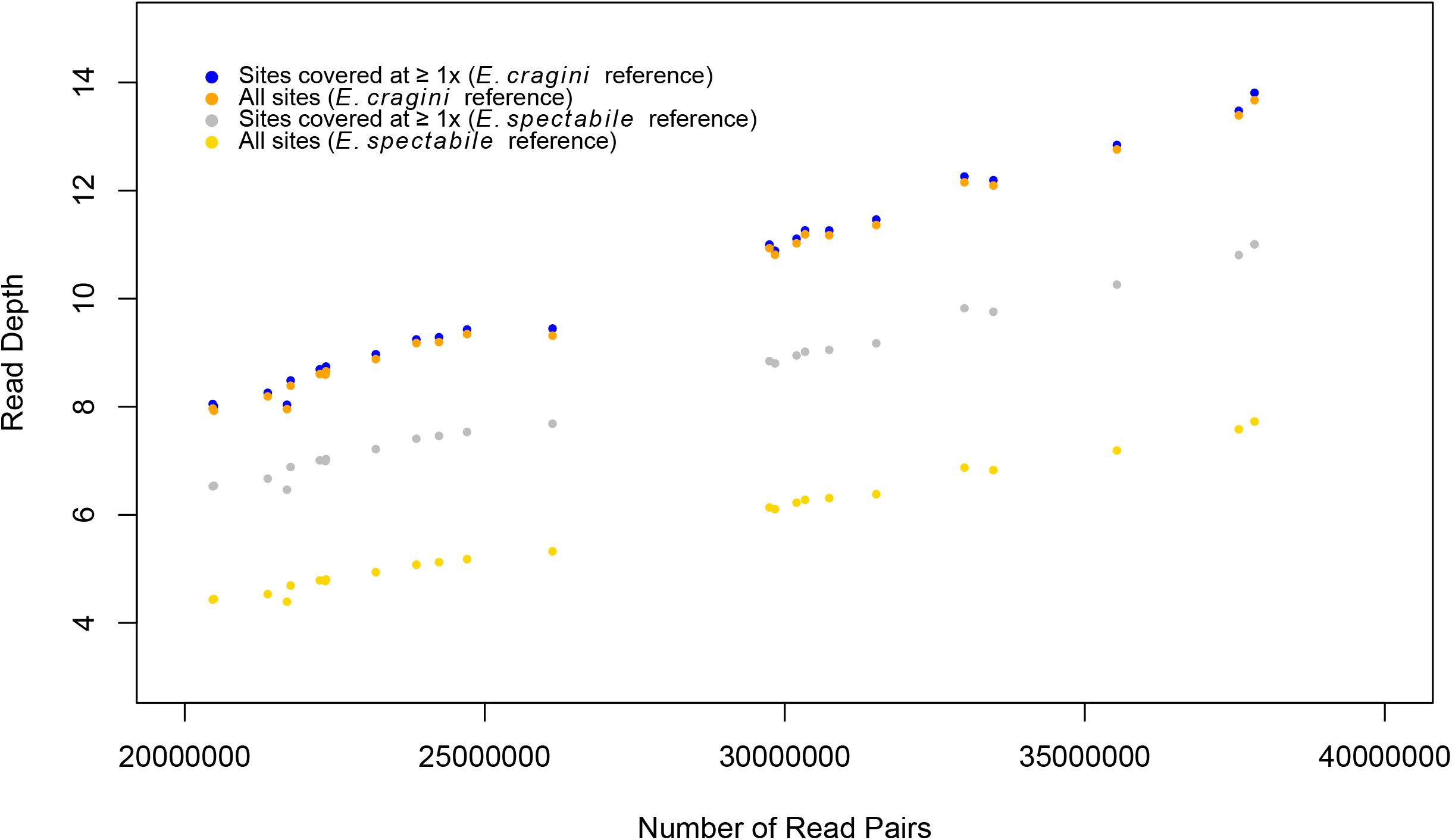
Read depth and number of read pairs for whole genome sequencing data. Average depth is shown for either a subset of sites with at least 1x coverage or for all sites in the genome, with alignment to either the conspecific *E. cragini* genome or the heterospecific genome of a closely related species (*E. spectabile*).

### Rapture quality control and coverage across species

Between 15.6% and 38.38% of total reads were identified as clones, and the proportion of reads identified as clones decreased in later Rapture batches (Supporting Figure 4). Most of the samples (96%) sequenced using Rapture generated >10,000 read pairs, and 93.7% generated >100,000 read pairs. For *E. cragini* samples with >10,000 read pairs, a high proportion of reads (generally >90%) mapped to the *E. cragini* reference genome. There were batch effects in proportion of reads mapping to the reference, with a higher proportion mapping in earlier Rapture batches. Approximately 90% of reads from *E. caeruleum* samples and approximately 80% of *E. spectabile* and *E. flabellare* reads mapped to *E. cragini* genome, with lower mapping success for these samples compared to *E. cragini* sequenced in the same batch (Supporting Figure 5a). The proportion of reads aligning to Rapture loci ranged from 46% to 70% and also displayed batch effects, as well as a decrease in reads mapping to Rapture loci with total read number (Supporting Figure 5a). Alignment of *E. caeruleum* and *E. spectabile* reads to the Rapture loci displayed similar patterns to *E. cragini* reads from the same batch, while the proportion of *E. flabellare* sequences aligning to the Rapture loci was distinctly lower compared to *E. cragini* from the same batch (Supporting Figure 5a). A lower proportion of *E. cragini* and *E. flabellare* sequences mapped to the heterospecific *E. spectabile* genome across batches, while a higher proportion of *E. caeruleum* and *E. spectabile* reads mapped to this reference genome (Supporting Figure 5b). The proportion of mapped reads aligning to the Rapture loci, however, was generally highly similar across species (Supporting Figure 5b).

The number of Rapture loci covered increased with number of reads for a given individual and tended to reach an asymptote above 10,000-100,000 reads (Figure 3, Supporting Figure 6). The maximum number of loci covered varied between species and between types of loci. For *E. cragini*, nearly all of the 2,119 Rapture loci were covered at each read depth. For the other species, a maximum of 1,700-1,800 of the Rapture loci were covered (Figure 3). The reduction in covered loci mainly came from a loss of short loci, of which only ~1,500 of 1,841 (~80%) were covered. A higher proportion of long loci (88%-95%) were sequenced at high coverage, and almost all of the outlier loci were sequenced at high coverage as well (Figure 3). Coverage for Rapture loci was nearly identical when the heterospecific *E. spectabile* reference genome was used for alignment (Supporting Figure 6). Per-base read depth was high for the portion of each locus covered by the capture baits for both long and short loci, representing large numbers of forward reads starting from the cut site overlapping the same region, although short loci had lower read depth beyond the capture baits compared to long loci (Supporting Figure 7).

**Figure 3.**
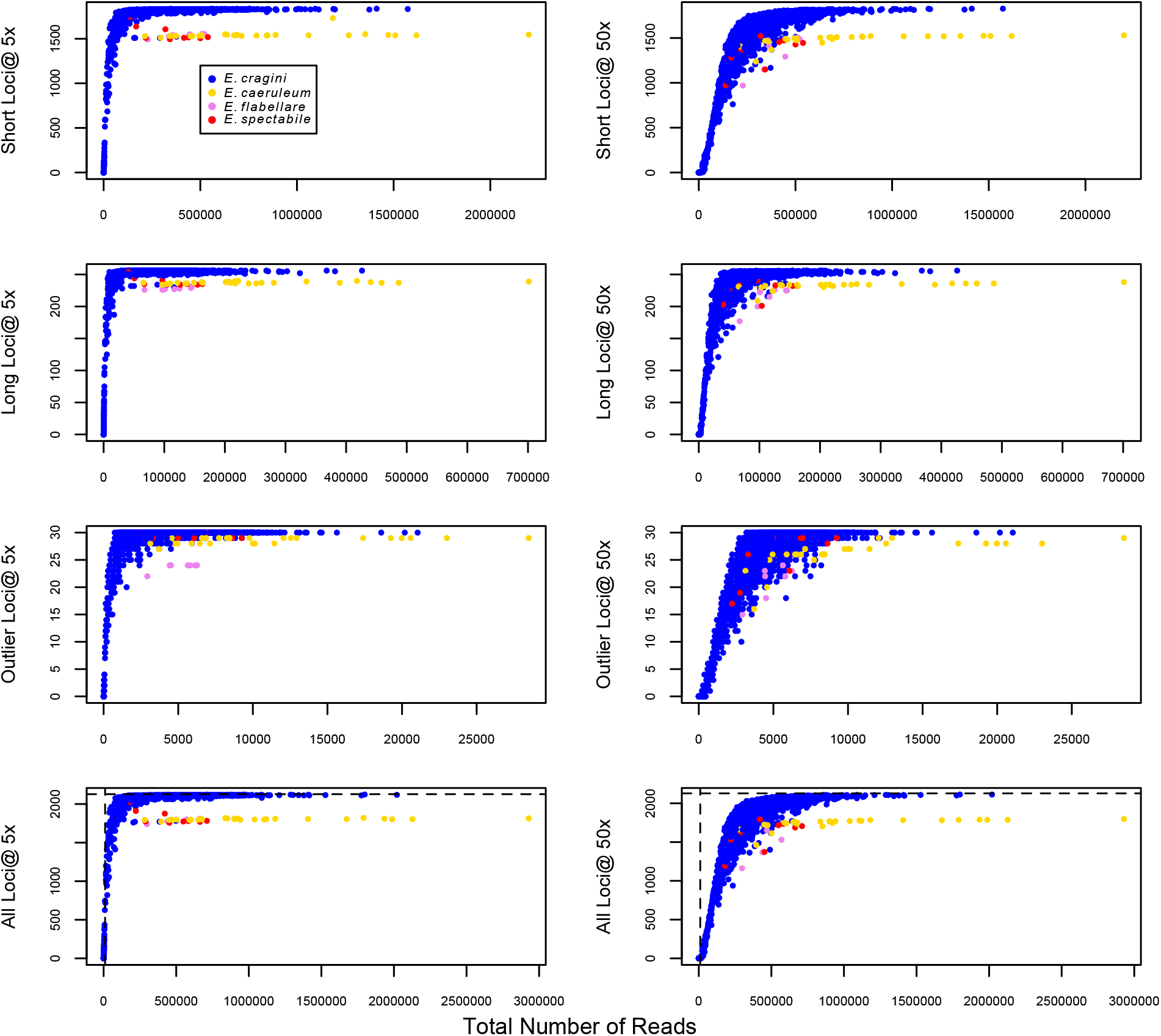
Coverage for Rapture loci (either all loci combined or short, long, and outlier loci taken separately) mapped to the *E. cragini* genome across four *Etheostoma* species.

### Polymorphism and heterozygosity

For the full Rapture dataset, there were 8,694 SNPs for the alignment to *E. cragini* and 10,495 SNPs for the alignment to *E. spectabile* across all 2,119 Rapture loci after filtering, indicating the presence of multiple SNPs per locus. The number of SNPs detected was similar for the subsetted Rapture datasets (8,581 SNPs for the alignment to *E. cragini* and 10,339 SNPs for the alignment to *E. spectabile*). For the WGS dataset, there were 5,759,437 SNPs for the alignment to *E. cragini* and 14,020,671 SNPs for the alignment to *E. spectabile*. Individual SNP heterozygosities were highly correlated across datasets – however, estimated heterozygosities were higher for the WGS datasets, and heterozygosity was higher for the *E. spectabile* WGS dataset than the *E. cragini* WGS dataset (Figure 4).

**Figure 4.**
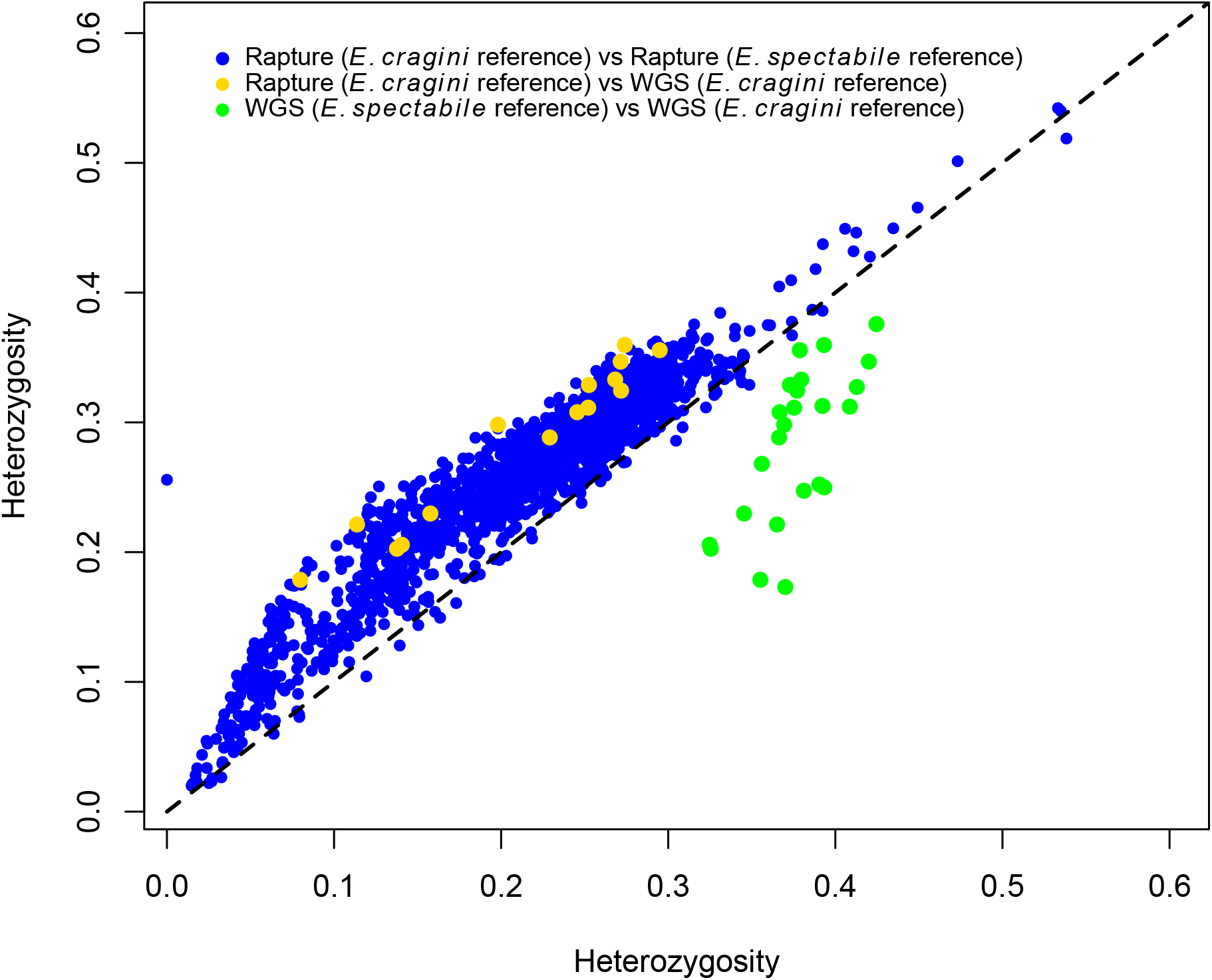
Comparisons of heterozygosity estimates across different datasets. For each comparison, the first estimate is plotted on the x-axis and the second is plotted on the y-axis. A 1:1 dotted line (expected for complete agreement across datasets) is also shown.

### Population structure and selection

We compared the results of population structure and selection in analyses for *E. cragini* among different datasets (full and subsetted Rapture datasets versus WGS) to evaluate how data type and reference genome affected downstream population genetic inferences. In contrast to batch effects on mapping and alignment to Rapture loci, we did not see strong evidence for batch effects in PCA-based analyses, and samples tended to cluster strongly by metapopulation rather than by batch (Supporting Figure 8). The admixture analysis in PCAngsd indicated that the best population delineation included 16 different populations for the both the full and subsetted Rapture datasets aligned to either reference genome (Figure 5a, Supporting Figure 9a-c). For WGS data, however, PCAngsd found 3 populations for the data aligned to the *E. cragini* reference, and 2 populations for the data aligned to the *E. spectabile* reference (Figure 5b, Supporting Figure 5d). The populations resolved for Rapture datasets broadly corresponded to major river drainages. The populations resolved for WGS lumped together populations in the major northern and southern drainages.

**Figure 5.**
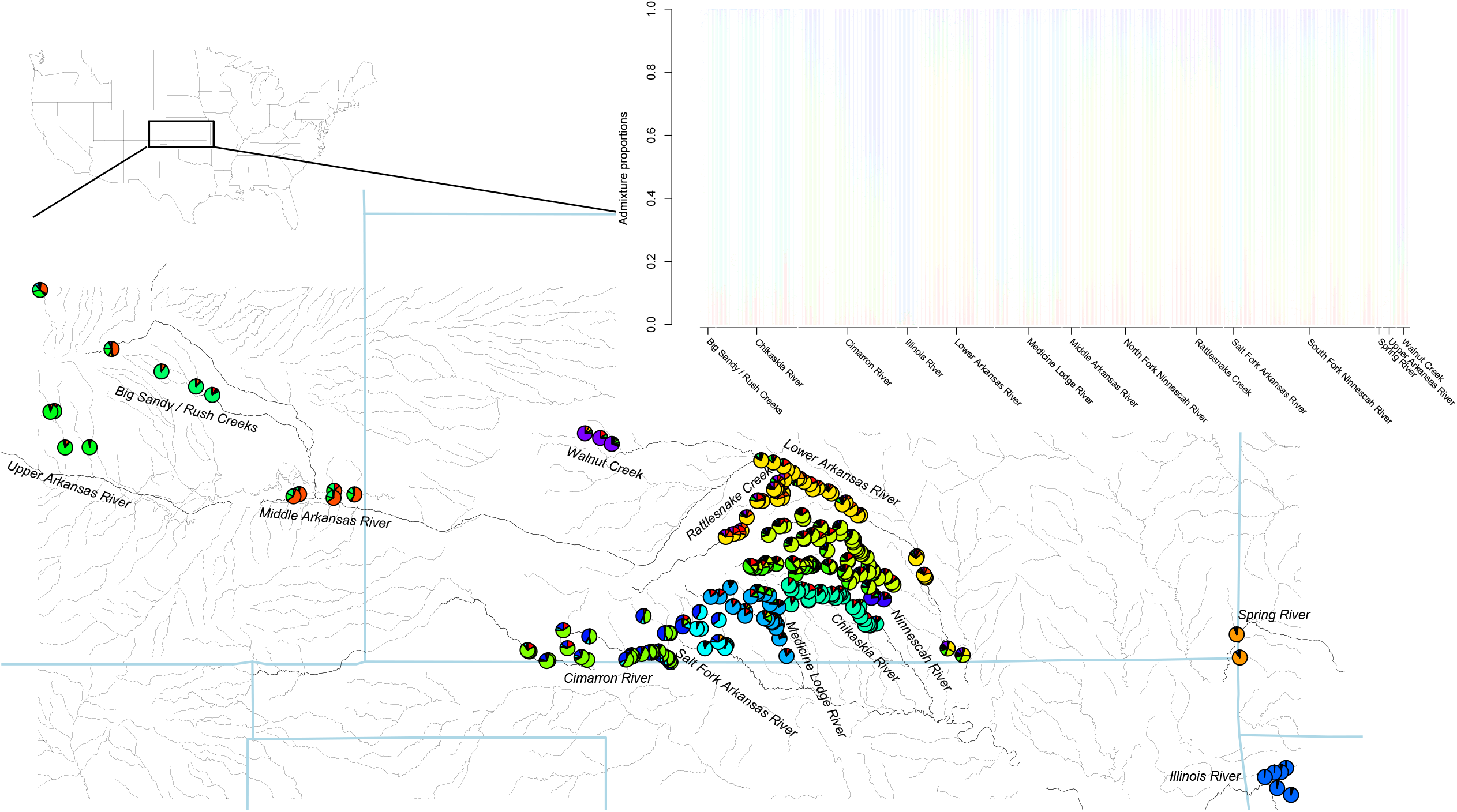
Admixture plots and mapped ancestry proportions. Each line in barplots represents an individual, and colors represent proportion of ancestry for each individual assigned to a given population. For the maps, pie charts represent either ancestry proportions aggregated for all individuals at a given site (for Rapture data) or admixture proportions for a single individual (for WGS data). Text on barplots indicates drainage of origin. Figure 5a. Rapture loci, full dataset, *E. cragini* reference.

**Figure 5b.**
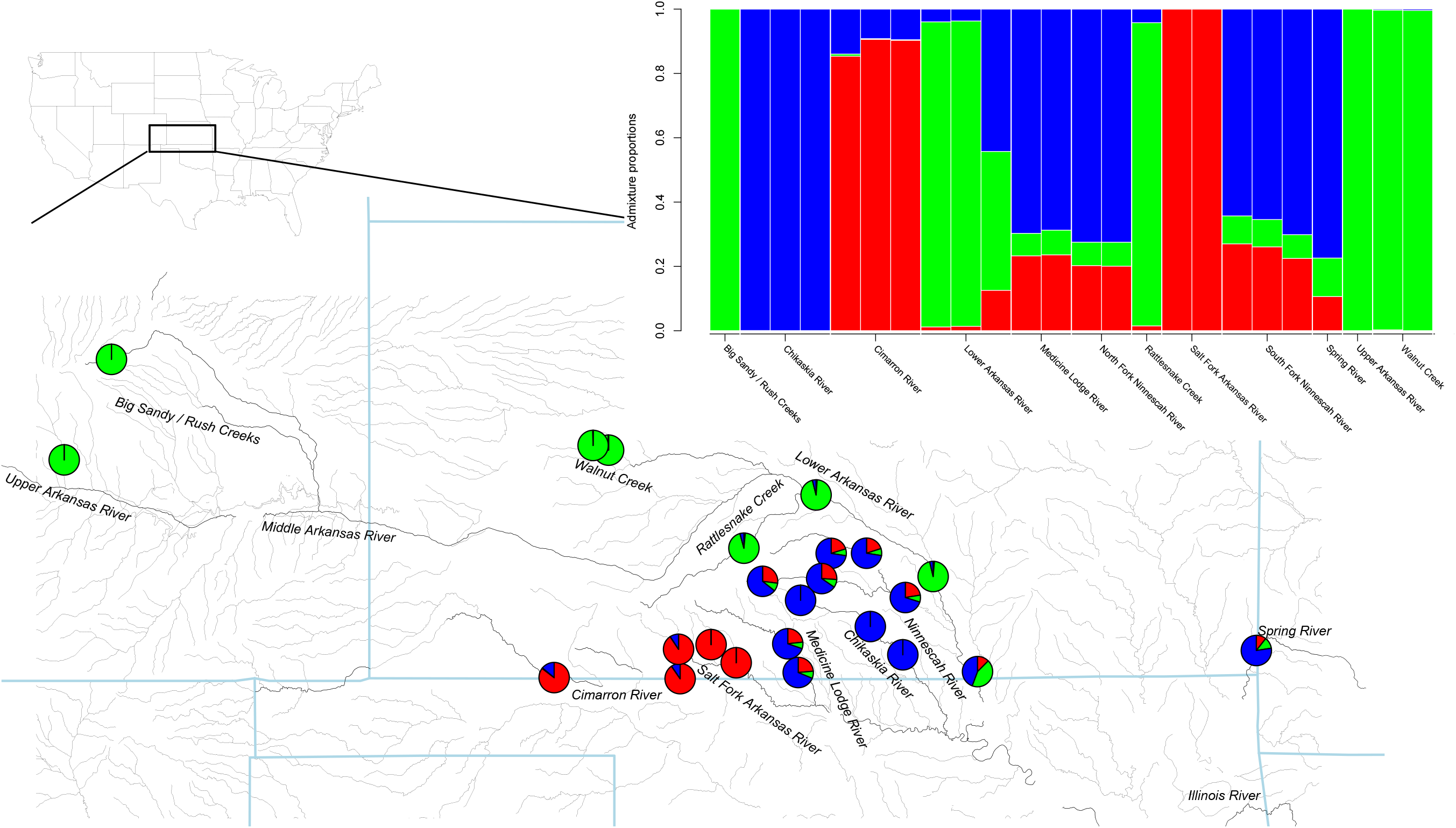
WGS data, *E. cragini* reference

For the both the Rapture and WGS datasets, PCAngsd did not identify any loci with significant evidence for selection when aligned to either reference genome after correction for the large number of tests (Supporting Figure 10).

### Phylogeny

Maximum likelihood phylogenetic analyses indicated that the Rapture loci were capable of resolving phylogenetic relationships with fairly strong support. The ML analysis produced 100% bootstrap support for correctly grouping *E. spectabile* with *E. caeruleum* and for grouping *E. flabellare* with *E. cragini*, as well as for grouping all individuals within their respective species (Figure 6a). Several deep phylogenetic splits (approximately 2.5-6 million years old) within *E. cragini* also received high support, and individuals within sites and within drainages were often grouped together with high support. Within *E. cragini*, populations showed a nested phylogeographic structure, with Arkansas populations basal to all populations to the east, and populations in east Kansas basal to populations further west. There was also a strongly supported split between populations in the mainstem Arkansas River and its tributaries and populations in drainages to the south of the mainstem Arkansas River (Figure 6b).

**Figure 6a.**
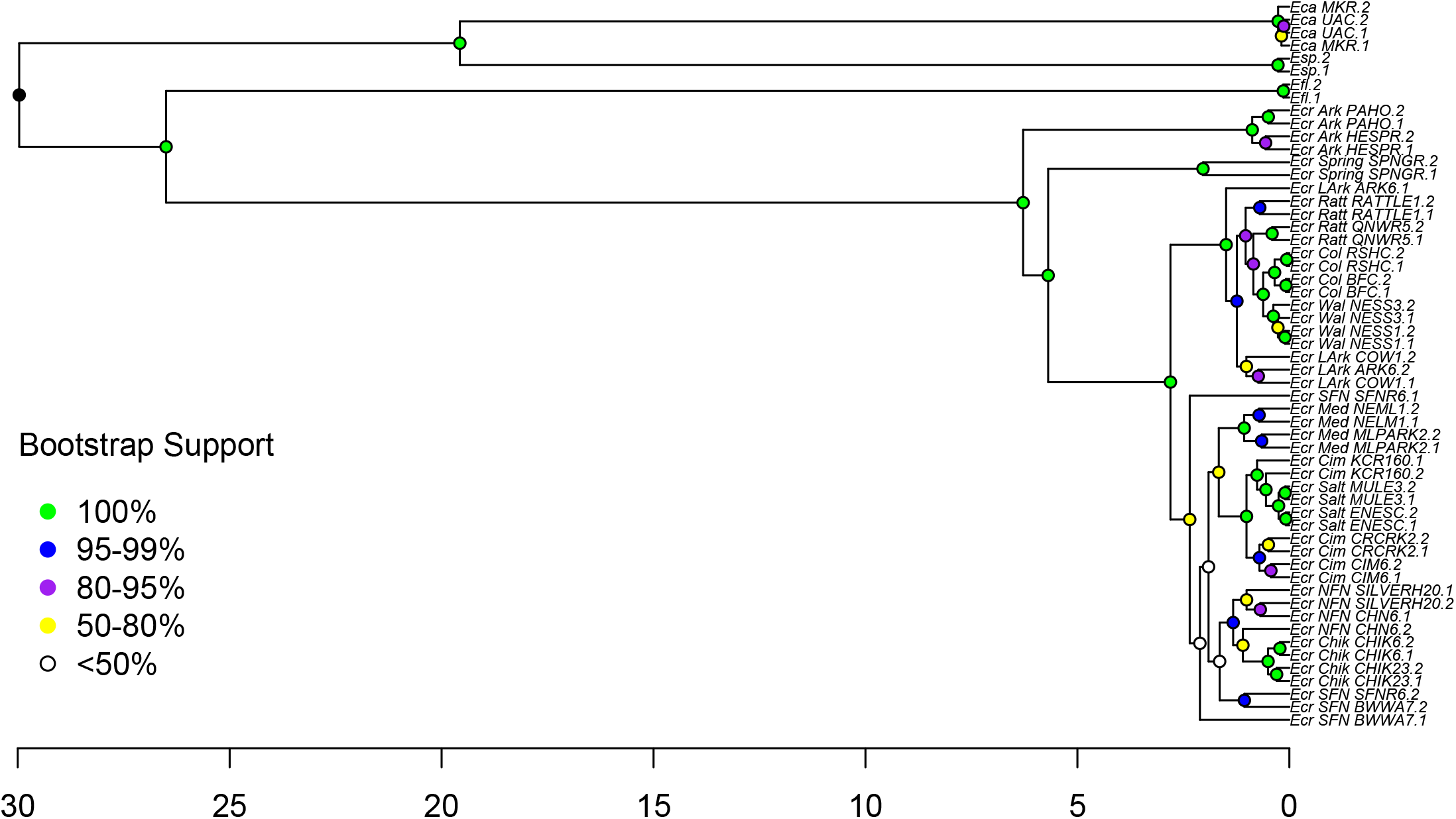
Time-calibrated maximum likelihood phylogeny for Rapture data. Eca = *E. caeruleum*, Esp = *E. spectabile*, Efl = *E. flabellare*, Ecr = *E. cragini*. Node labels for *E. caeruleum* individuals include a site identifier, and node labels for *E. cragini* individuals include a metapopulation identifier followed by a site identifier. Time (on the x-axis) is expressed in millions of years ago.

**Figure 6b.**
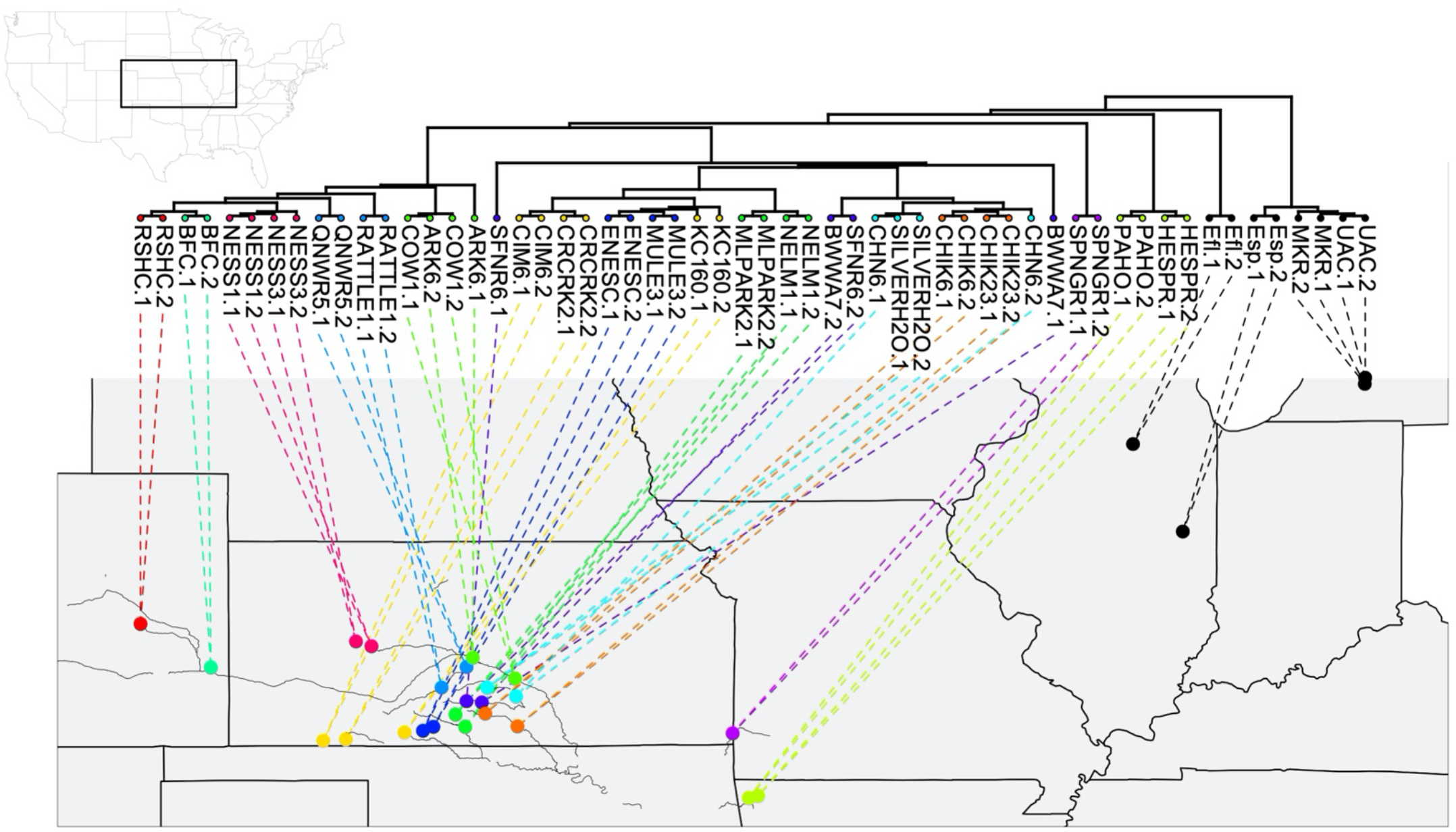
Phylogeny plotted in space.

Per-site phylogenetic informativeness profiles for the three categories of loci showed similar overall patterns from 30 million years ago to 2-3 million years ago, with a slightly convex but relatively stable informativeness profile over time (Supporting Figure 11). Informativeness dropped rapidly from 2 million years ago to the present for the long and short loci, but outlier loci exhibited a secondary peak from 1-2 million years ago for the outlier loci followed by a steep decline. Long loci tended to have lower per-site phylogenetic informativeness than short or outlier loci.

## Discussion

There are a number of common questions any researcher involved in the design and implementation of a population genomic or phylogenomic study in a non-model organism will have to address. These include: how many loci and how many individuals do I need to include? Should I sequence loci over the entire genome or should I use sequence capture to target a smaller number of loci at high depth? Should I generate a reference genome for my species or will I be able to use a reference genome from a closely related species, and how will this choice affect the interpretation of my data? Will one methodology work equally well across all target populations and species? And how cost-effective are these alternative methods? All of these questions are perhaps even more relevant for projects aimed at diverse species radiations, as such projects by their nature encompass a number of closely related species. Based on the work described here, we discuss how these questions can be addressed and which methods are most appropriate for different applications.

### To Rapture or not to Rapture (and how to Rapture)

A number of sequence capture methods exist, ranging from anchored probes (Lemmon et al. 2012) and ultraconserved elements (Faircloth et al. 2012) developed for use across a wide variety of taxa, to more focused methods that develop and use a bait set for a single species (Margres et al. 2018). Previous work with the Rapture method in marine turtles demonstrated that baits developed for a single species work well in related species that diverged tens of millions of years ago (Komoroske et al. 2019), and we confirm in this work that Rapture loci developed for a single darter species can also be used in other species from the same group. Rapture loci were recovered with highest coverage from the target species (*E. cragini*) but a majority of loci were recovered from all four species. Long loci spanning both sides of the restriction site were recovered with higher frequency than loci that did not span the restriction site (short loci) across species, and we obtained higher coverage in regions flanking the RAD locus for long loci as well. This is possibly because of a greater possibility of bait capture for more dissimilar sequences with more baits per locus (5 baits for long loci compared to 2 baits for short loci). We also used 80-bp tiled baits as opposed to 120-bp baits used in previous Rapture studies, which may have improved the likelihood of capture as well. These results suggest overall that using loci that span both sides of the restriction site and using 80-bp tiled baits will likely lead to the most consistent recovery of Rapture loci across related species. Incorporating multiple reference genomes or creating pseudo-reference genomes from pilot RAD data for other species of interest may be useful in designing bait sets that will function best across species radiations.

One of the goals of a Rapture approach is to consistently genotype a large number of loci for a large number of samples. Processing a very large number of samples will necessitate splitting these samples into batches, and batch effects are known to plague some HTS analyses (Leigh et al. 2018, Lambert et al. 2019). The genotyping described here was conducted in 5 batches over a time period of approximately 8 months, and we did find evidence for batch effects in some aspects of Rapture data generation and analysis, specifically in the proportion of clonal reads and the proportion of reads mapping to reference genomes and Rapture loci. The decrease in clonal reads likely resulted from using fewer PCR cycles in the final amplification step in later batches after we had determined that fewer cycles were needed to produce an adequate DNA concentration. Batch effects related to reference genome mapping could potentially be a downstream effect of the change in clone frequency, or they could also be related to somewhat reduced efficacy of the capture reaction over time (possibly due to freezing and thawing of reagents over time, although baits were aliquoted to minimize freeze-thaw cycles). PCA results, however, suggest that these batch effects did not have much downstream effect on the interpretation of the data, as on PC axes explaining most of the variation in the data samples did not cluster by batch, and rather clustered strongly by metapopulation. Additionally, high overall coverage across Rapture loci likely alleviates problems of nonrandom data missingness across batches and allelic dropout commonly seen in traditional RADseq data (Malinsky et al. 2018). Our conclusion is that batch effects should not strongly affect downstream analyses of Rapture data collected over multiple lab preparations and sequencing lanes.

### Conspecific vs heterospecific reference genomes

For capture-based reduced-representation genomic methods, the choice of reference genome impacts both the initial phase of SNP discovery and development of capture baits as well as the analysis phase. Due to the currently limited availability of reference genomes and the time and cost required to sequence a novel genome, researchers initiating a sequence capture project may need to use a heterospecific genome for the initial SNP discovery and bait design steps, as we did here. Studies in birds have suggested that heterospecific reference genomes can be useful for SNP discovery (Galla et al. 2019), although strongly conserved genome structure across bird taxa (Ellegren 2010) may increase the utility of heterospecific reference genomes for SNP discovery in this group. In this study, using a heterospecific reference genome for the initial design phase resulted in baits that successfully captured polymorphic RAD loci in our target species (*E. cragini*) and in congeneric species. This may reflect conserved chromosome number and large regions of synteny among genomes for species belonging to this group (Supporting Information 1). However, comparison of the *E. cragini* and *E. spectabile* genomes indicated substantial changes in genome size and organization among species within *Etheostoma*, suggesting that genome structure has evolved substantially over the approximately 30 million-year history of the genus. Genome structure and karyotype may in some cases vary widely within species radiations (e.g. Vershinina and Lukhtanov 2016), and as such caution should still be exercised when using heterospecific reference genomes in SNP discovery and bait design. As more eukaryotic genomes become available, adopting a pan-genome approach, used in the past to identify core regions common to prokaryotic genomes within specified taxonomic groups (Vernikos et al. 2015)), could become an attractive alternative to using a single reference genome. This approach may be particularly appealing to researchers working with species radiations, as targeting genomic regions that are conserved throughout a radiation should increase the utility of the capture bait set across species.

Choice of reference genome will also impact the downstream analysis and interpretation of both targeted sequence capture and WGS data. Although mapping sequence reads generated from one species to the genome of a closely related species is still common practice, the effects of mapping reads to a heterospecific genome versus a conspecific genome are still relatively understudied. Galla et al. (2019) mapped RADseq and low-coverage WGS data to either a conspecific, congeneric, confamilial, or conordinal genome and found a decreasing alignment rate with increasing phylogenetic distance, as well as less consistency in estimates of genetic diversity when reads were mapped to a more distantly-related genome. Our WGS results generally agree with these findings. Mapping reads generated from low-coverage WGS of *E. cragini* individuals to the *E. cragini* reference genome was generally more successful than mapping to the *E. spectabile* genome. Lower read depth and allelic dropout could contribute to different estimates of genetic diversity from conspecific and heterospecific reference genomes. However, even though we inferred population structure using genotype likelihoods, which should mitigate the effects of lower read depth associated with mapping to a heterospecific reference genome (Nevado et al. 2014), admixture results were also affected by the choice of reference genome. The existence of multiple rearrangements among darter genomes observed in this study and others (Moran et al. 2020) could aggravate the effects of using a heterospecific reference genome. For our Rapture dataset, however, the effects of mapping to a heterospecific reference were much reduced, and downstream inferences regarding diversity and population structure were similar, regardless of which reference genome was used for mapping. This suggests that sequence capture may reduce biases associated with the absence of a closely-related reference genome, possibly because the RAD loci targeted by sequence capture in this case were designed by alignment to a heterospecific reference genome and thus were fairly conserved across genomes.

### Effects of reference genome choice and data type on population genomic inferences

Different data types can be differentially suited to different analyses. We found that Rapture was much better at identifying fine-scale population structure than WGS. This is likely partially due to the much greater spatial coverage and the greater number of individuals we were able to sequence via this method with a similar amount of sequencing effort. Higher coverage overall for the Rapture data may also alleviate allelic dropout and decreased sensitivity for calling heterozygotes associated with low-coverage WGS. However, as PCAngsd uses genotype likelihoods rather than called genotypes, this issue may not have strongly affected these analyses.

Previous work has asserted that WGS is typically better suited to detecting evidence of selection using genome scan methods than RAD-based approaches, which target relatively small portions of the genome (Lowry et al. 2016). For selection scan methods to be accurate, however, they must take into account variation in allele frequencies due to neutral processes, such as change in population size over time or spatial population structure (de Villemereuil et al. 2014). This may be particularly important in species with limited dispersal capability, such as freshwater fish (Shurin et al. 2009). We found little evidence for selection in any of the Rapture datasets, even for loci that showed some evidence of selection in the pilot dataset. As the pilot dataset did not delineate many of the fine-scale populations indicated by the full or subsetted Rapture datasets, however, we believe it is very plausible that the outlier loci identified in these preliminary analyses represent loci that were differentiated due to neutral population within broad-scale groupings rather than selection. We found little evidence of selection for the WGS datasets as well. This may be due again to the lower number of individuals sampled for WGS, and less accurate estimation of population structure may also have confounded detection of loci under selection using WGS. Future analyses of selection using low-coverage WGS data may also benefit from the application of methods that estimate linkage disequilibrium based on genotype likelihoods and prune closely linked SNP loci (Fox et al. 2019), which would reduce overall SNP density but also potentially reduce false positives and increase power by lowering thresholds for accurately detecting loci under selection while accounting for multiple comparisons.

RADseq-based methods can detect selection if marker density is high relative to the size of linkage disequilibrium blocks (Catchen et al. 2017), and Rapture workflows designed to detect selection with these factors in mind may be comparable to WGS. Alternatively, Rapture methods can also include loci with known *a priori* effects on fitness (such as loci associated with disease susceptibility). While our Rapture loci identified *a priori* as under selection by genome scans in the pilot dataset did not show strong evidence of selection in the larger datasets, Rapture panels designed to include high marker density as well as immune-associated loci constituted an effective means identifying loci associated with survival in female Tasmanian devils (*Sarcophilus harrisii*) with a transmissible cancer (Margres et al. 2018).

### Phylogenetic informativeness of Rapture loci

Sequence capture strategies targeting ultra-conserved elements (UCEs) and protein-coding genes have been evaluated in the past for percomorph fishes (Gilbert et al. 2015). UCE flanks and protein-coding genes in general showed great utility for resolving deeper split but a loss of phylogenetic signal for more recent epochs, with per-locus phylogenetic informativeness for UCE flanks and protein-coding genes peaking between 20-40 million years ago and exhibiting rapid decline from 20 million years ago to the present. The phylogenetic informativeness of Rapture loci was fairly constant over time and potentially more useful for examining relatively recent splits between closely related species. However, phylogenetic informativeness for Rapture declined rapidly for very recent epochs. This is also potentially reflected in support values estimated here for relationships within the *E. cragini*. We obtained 100% bootstrap support for older splits between populations in Arkansas versus populations further east, as well as high support for a sister relationship between populations in eastern Kansas and all other populations to the west and a split dating to approximately 3 million years ago between populations in drainages associated with the mainstem Arkansas river and populations in drainages to the south of the Arkansas River. For more recent splits, support values were overall fairly high (95-100%) but much lower for some nodes, indicating ambiguous support for some relationships. This likely represents both a true lack of phylogenetic informativeness (i.e. substation rates too low to allow for reliably distinguishing among alternative relationships) as well as potentially other confounding factors, such as gene flow and maintenance of ancestral polymorphisms. Alternative methods that incorporate gene flow and demographic modeling (Jackson et al. 2017, Scott et al. 2018) could allow for more reliable inferences for recently diverged populations.

### Costs and benefits of different sequencing methods

With limited funding, cost will always be a consideration. Rapture has a somewhat costly initial investment but is still highly cost-effective ($13.42 sample including bait design and production for 1,900 individuals in our study, or <$10 per sample if baits are already available; Table 1) in terms of cost per sample when compared to either BestRAD or low-coverage WGS. Given these low costs, Rapture is a very attractive method for conducting future work in the darter system, especially when extensive individual-level and spatial sampling are important components of the project design. The data produced by Rapture can be supplemented by low-coverage WGS if this is needed for the study, and the relatively high cost per individual of WGS in this study (~$275 per sample) could potentially be reduced by using poolseq (Schlötterer et al. 2014).

Overall, the Rapture method outlined here represents a potentially powerful methodology for phylogenomics and population genomics, both in darters and in diverse radiations of non-model organisms more generally. We have also shown several potential pitfalls associated with using heterospecific genomes. While targeted sequence capture seems to mitigate some of these pitfalls, choosing to use a heterospecific reference genome still has consequences that should be carefully considered during study design. As more reference genomes and sequence capture methods become available, Rapture will become an increasingly attractive option, especially when large sample sizes, extensive spatial coverage, or high read depth are important.

## Supporting information

Supporting Table 1 (sample metadata)

## Acknowledgments

This work was funded by grant #SCTF911C from the Colorado Parks and Wildlife Species Conservation Trust Fund and grant # E-30-R-1 from the Kansas Department of Wildlife, Parks, and Tourism. We would like to thank Harry Crockett (Colorado Parks and Wildlife Agency), Jordan Hoffmeier and Mark van Scoyoc (Kansas Department of Wildlife, Parks, and Tourism), and Brian Wagner (Arkansas Game and Fish Commission) for providing samples. Rainbow darters were collected under a scientific collector’s permit issued by the Michigan Department of Natural Resources, and collection of *E. spectabile* and *E. flabellare* was approved by the Illinois Department of Natural Resources under Scientific Collecting Permit A15.4035.

We thank M. Meek and D. Oliveira for providing helpful feedback on our manuscript. This is W.K. Kellogg Biological Station publication number xx.

## Data accessibility statement

The *E. cragini* Whole Genome Shotgun project has been deposited at DDBJ/ENA/GenBank under the accession JAAVJE000000000. The version described in this paper is version JAAVJE010000000. Short-read data have been uploaded to the NCBI as BioProject PRJNA611833. Analysis scripts and capture bait sequences are available at https://github.com/nerdbrained/darter_rapture.

## Notes

https://github.com/nerdbrained/darter_rapture

https://www.ncbi.nlm.nih.gov/bioproject/611833

### Competing Interest Statement

The authors have declared no competing interest.

### Summary of Updates

We have revised this manuscript to include more quality control data (proportion clonal reads and proportion of reads aligning to reference genomes and Rapture loci), tests for batch effects, and additional filtering criteria for SNPs, which changed the results of downstream analyses somewhat. We have also included more discussion on the effects of reference genome choice on both the bait development and data interpretation steps.

