## Supporting Information, Figures, and Tables for "Rapture-ready darters: choice of reference genome and genotyping method (whole-genome or sequence capture) influence population genomic inference in *Etheostoma*"

**Supporting Information 1.**

We generated assembly statistics with QUAST v4.3 (Gurevich et al. 2013) to evaluate the quality of the *E. cragini* genome compared to the congeneric *E. spectabile* genome (v. UIUC_Espe_1.0, downloaded from NCBI; Moran et al. 2020) and to the confamilial *Perca flavescens* (v. PFLA_1.0, downloaded from NCBI; Feron et al. 2020), *Perca fluviatalis* (v. GENO_Pfluv_1.0; Ozerov et al. 2018), and *Sander lucioperca* (v. SLUC_FBN_1, downloaded from NCBI; Nguinkal et al. 2019) genomes. To assess completeness of the *E. cragini* assembly, we determined the number of Actinopterygii-specific Benchmarking Universal Single-Copy Orthologs (BUSCOs) identified with BUSCO v2.0 (Simão et al. 2015) using the actinopterygii_odb10 lineage dataset. We also used RepeatModeler (Smit et al. 2015) v.1.0.11 and RepeatMasker v.4.0.5 (Smit and Hubley 2015) to identify repetitive elements in the *E. cragini* assembly and in the most recent version of the *P. fluviatalis* genome. We combined this information with published data to compare the proportion of repetitive elements among percid fish genomes. To compare genomic synteny and homology among genomes, we aligned the Dovetail sequence with confamilial genomes that contained chromosome-level assemblies (*E. spectabile*, *P. flavescens*, and *P. fluviatalis*) with Minimap2 (Li 2018) using the D-Genies web interface (dgenies.toulouse.inra.fr; Cabanettes and Klopp 2018). We visualized the alignment in D-Genies with a dotplot, using the “strong precision” setting to remove small matches. We converted the annotations for the *E. spectabile* genome into ENSEMBL format using a script (<https://github.com/NBISweden/EMBLmyGFF3>) and transferred annotations from the *E. spectabil*e genome to the *E. cragini* genome using RATT v.0.95 (Otto et al. 2011).

The *E. cragini* reference genome generated by Dovetail included data from 461,647,761 2x150bp shotgun read pairs for the initial *de novo* genome assembly and approximately 194 million and 228 million 2x150bp read pairs for the Chicago and HiC libraries, respectively. The final assembly had an estimated physical coverage of 16,456.85x across 4,667 scaffolds, an L50 of 11, an N50 of 27.59 Mb, and a total length of 643.1 Mb (Supporting Table 2). A total of 97.6% of the genome was assembled into 24 chromosomes. The final *E. cragini* assembly size was smaller than predicted by initial estimates based on short-reads (31-mer estimated size of = 736.9 Mb), and was also smaller than all other percid genomes sequenced to date. However, the number of complete BUSCOs observed in the *E. cragini* assembly was comparable to previously published percid assemblies (Supporting Table 2), despite being 68-75% of the size of these other assemblies. Within the *E. cragini* assembly, 172 Mb (26.7% of total genome size) was classified as repetitive, representing both a lower total amount of repetitive sequence and a lower proportion of the genome classified as repetitive compared to other percids (Supporting Table 3).

Genome alignments showed many long regions of synteny between the two genomes, with 24 long scaffolds of the *E. cragini* genome mostly aligning to single chromosomes in the published genomes (Supporting Figure 1). There was also extensive evidence, however, for inversions and chromosomal rearrangements in all 24 chromosomes between the species, often near the ends of chromosomes. Inversions and rearrangements were more pronounced in the *E. cragini – E. spectabile* alignment, and relatively few sequences from the *E. cragini* assembly aligned to the unplaced scaffolds in the *E. spectabile* assembly compared to the chromosome-level scaffolds, although the *E. cragini* – *E. spectabile* alignment had higher identity overall (Supporting Figure 3). RATT successfully transferred 99% of annotation elements (12,497 of 12,849 possible elements) across genomes, although some elements (14%) that were contiguous on the *E. spectabile* genome were split among several genomic regions or only partially transferred onto the *E. cragini* genome. All 740 gene models from the *E. spectabile* genome were transferred to the *E. cragini* genome, although 147 gene models (80%) were partially transferred with 161 exons found in the *E. spectabile* genome not found in *E. cragini*.

**Supporting Table 2.** Number (%) of complete (C), single-copy (S), duplicated (D), fragmented (F), and missing (M) Benchmarking Universal Single-Copy Orthologs (BUSCOs) out of 3,640 total in the Actinopterygii-specific set.

|  | Complete | Single-copy | Duplicated | Fragmented | Missing | Total |
| --- | --- | --- | --- | --- | --- | --- |
| *E. cragini* | 3449 (94.8%) | 3425 (94.1%) | 24 (0.7%) | 46 (1.3%) | 145 (3.9%) | 3640 |
| *E. spectabile* | 3442 (94.6%) | 3347 (92%) | 95 (2.6%) | 18 (0.5%) | 180 (4.9%) | 3640 |
| *P. flavescens* | 3539 (97.2%) | 3506 (96.3%) | 33 (0.9%) | 8 (0.2%) | 93 (2.6%) | 3640 |
| *P. fluviatilis* | 3509 (96.4%) | 3477 (95.5%) | 32 (0.9%) | 16 (0.4%) | 115 (3.2%) | 3640 |
| *S. lucioperca* | 3537 (97.1%) | 3499 (96.1%) | 38 (1%) | 11 (0.3%) | 92 (2.6%) | 3640 |

**Supporting Table 3.** Assembly statistics for the *E. cragini* assembly presented here compared to other recently released percid fish assemblies.

| **Species** | **Total Length** | **N50** | **L50** | **# Scaffolds** | **% Gaps** | **%GC** | **% Repetitive** |
| --- | --- | --- | --- | --- | --- | --- | --- |
| *E. cragini* | 643,100,042 | 27,593,271 | 11 | 4,667 | 0.520 | 40.52 | 26.7 |
| *E. spectabile* | 854,790,067 | 30,497,795 | 13 | 3,118 | 0.470 | 40.91 | 30.9 |
| *P. flavescens* | 877,456,336 | 37,412,490 | 11 | 268 | 0.047 | 40.84 | 41.7 |
| *P. fluviatalis* | 951,362,726 | 39,550,354 | 11 | 304 | 0.031 | 40.90 | 40.6 |
| *S. lucioperca* | 900,477,756 | 4,929,547 | 52 | 1,313 | 0.003 | 40.92 | 39 |

Supporting Figure 1. Preliminary sample sites and principal coordinate analysis

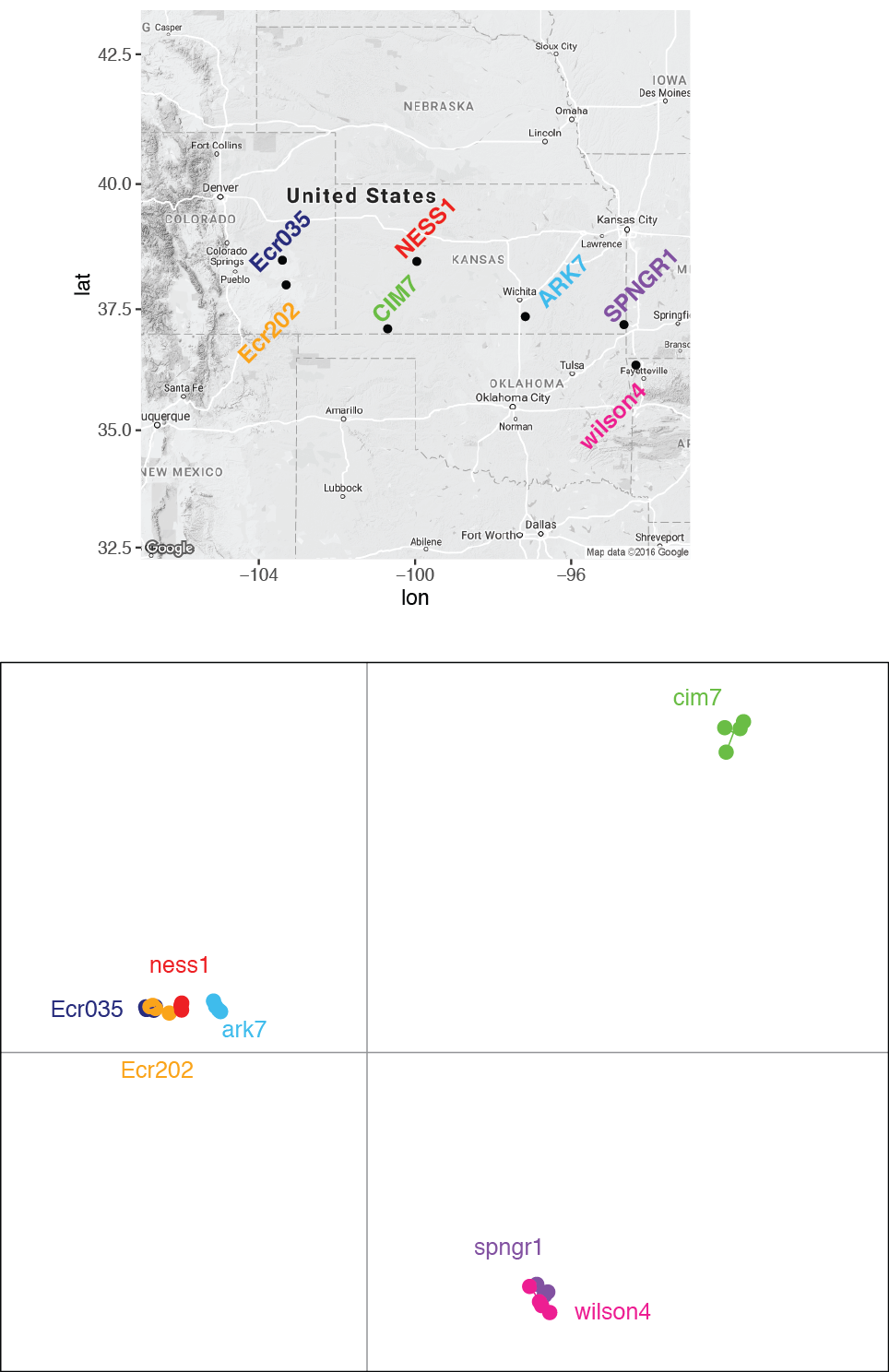

Supporting Figure 2. Bayescan results from pilot samples showing distribution of FST values across loci, with loci showing evidence of selection to the right of the vertical line. Top panel: rangewide analysis, bottom panel: mainstem Arkansas River population only.

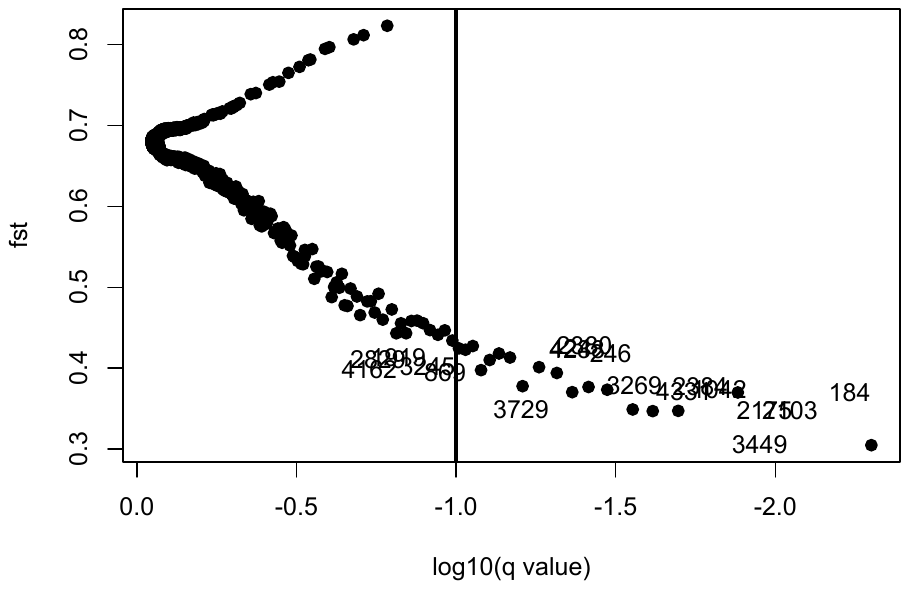

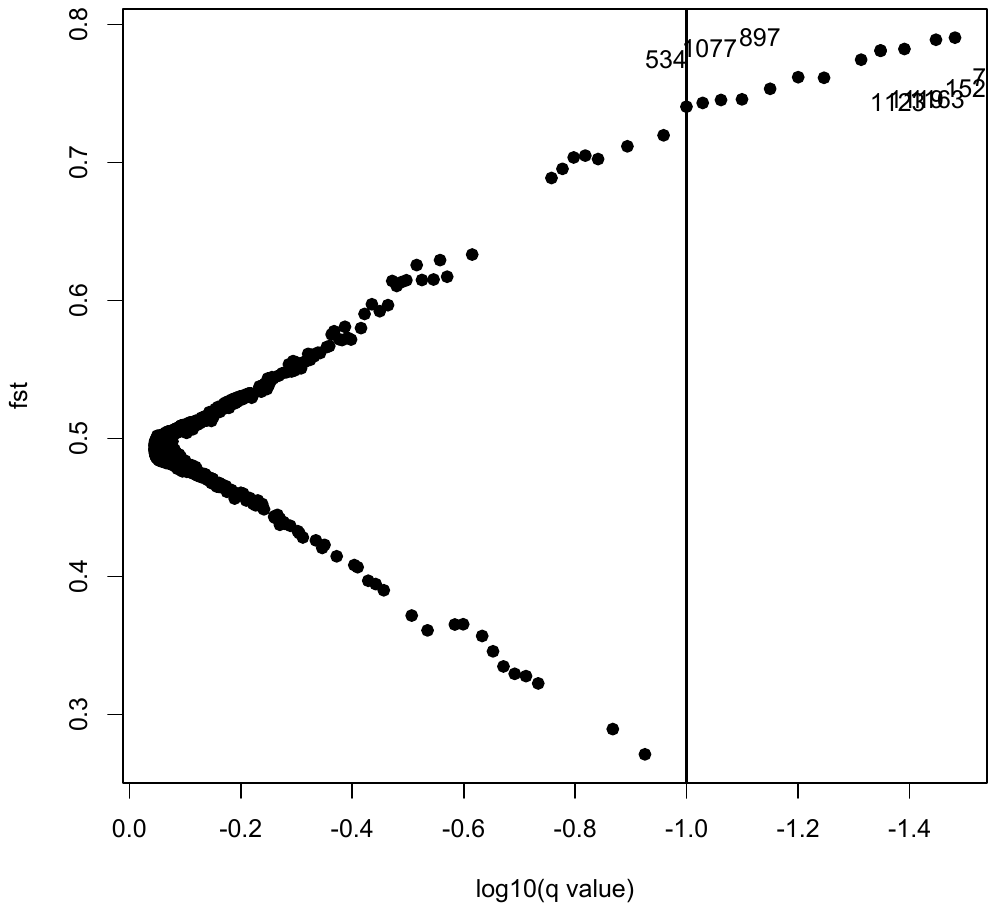

**Supporting Figure 3**. Dotplots of whole-genome alignments between *E. cragini* and other chromosome-level percid fish genome assemblies after removing weak-precision alignments. The *E. cragini* genome is shown on the y-axis in each case. A summary of the proportion of total matches at various levels identity is shown below each dotplot.

(a) *E. spectabile* vs. *E. cragini*.

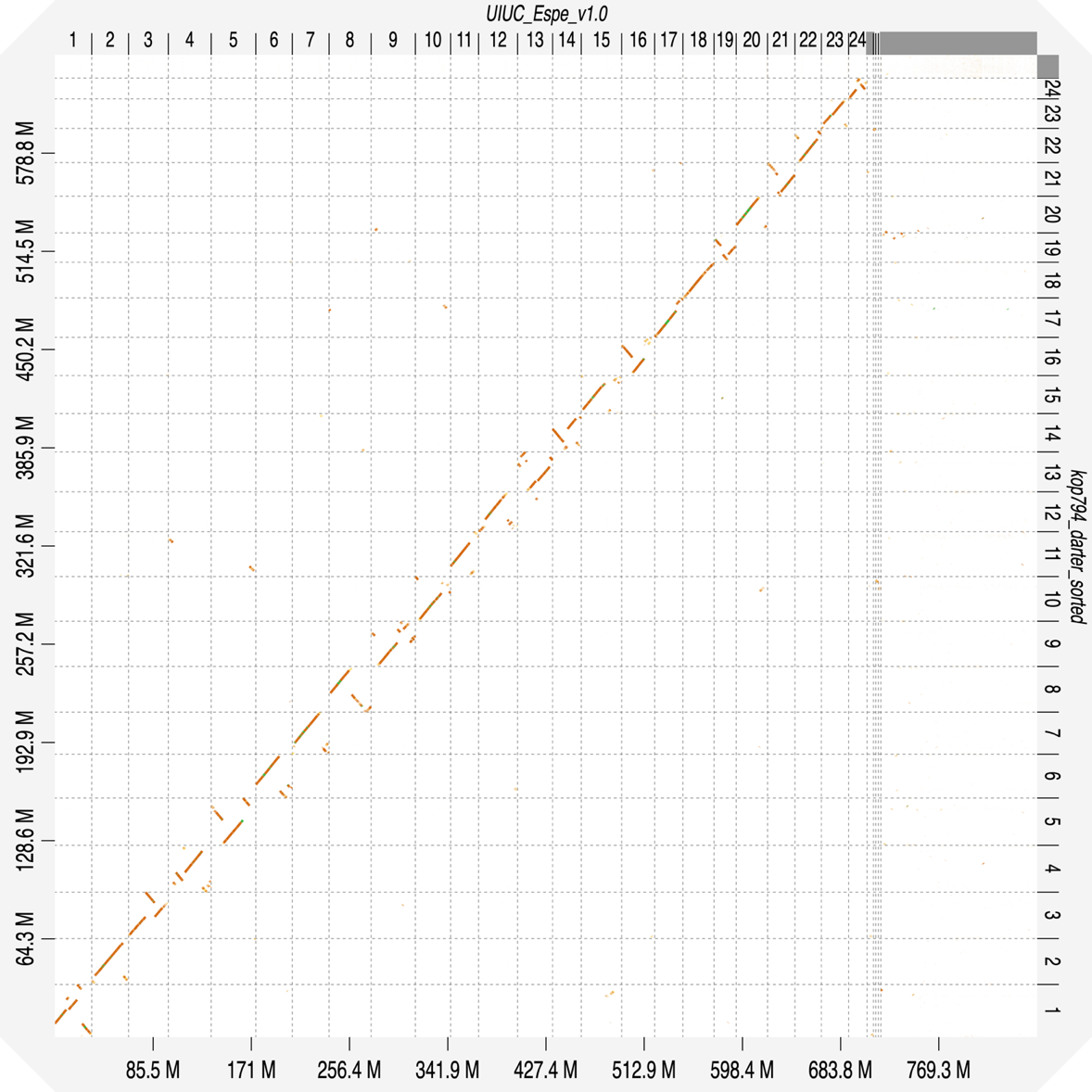

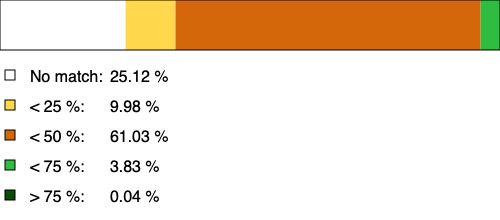

(b) *P. flavescens* vs. *E. cragini*.
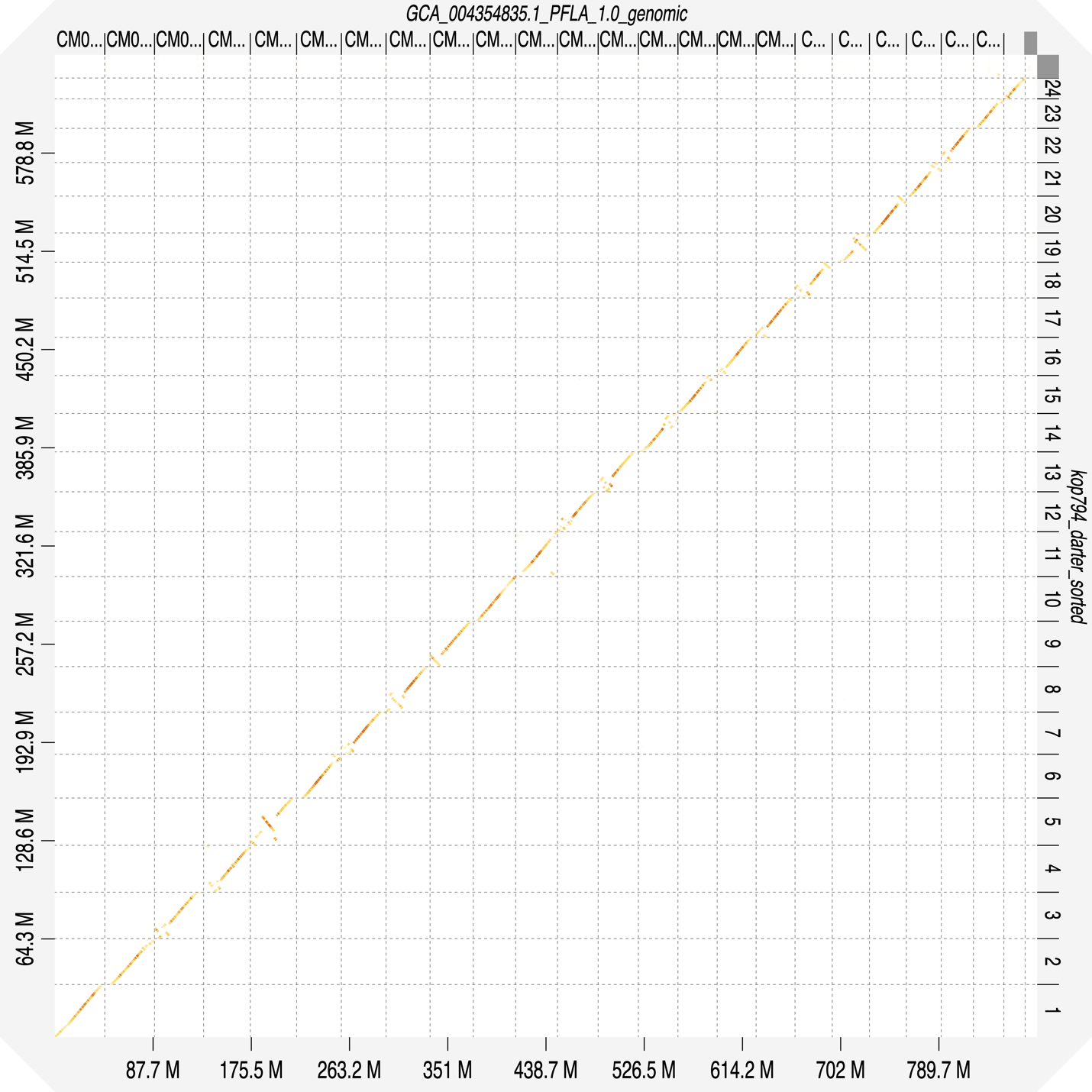

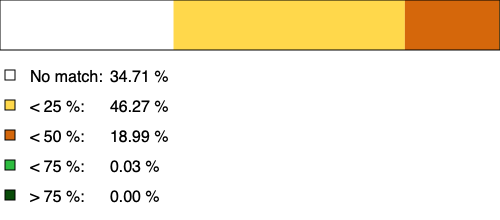

(c) *P. fluviatalis* vs. *E. cragini*.
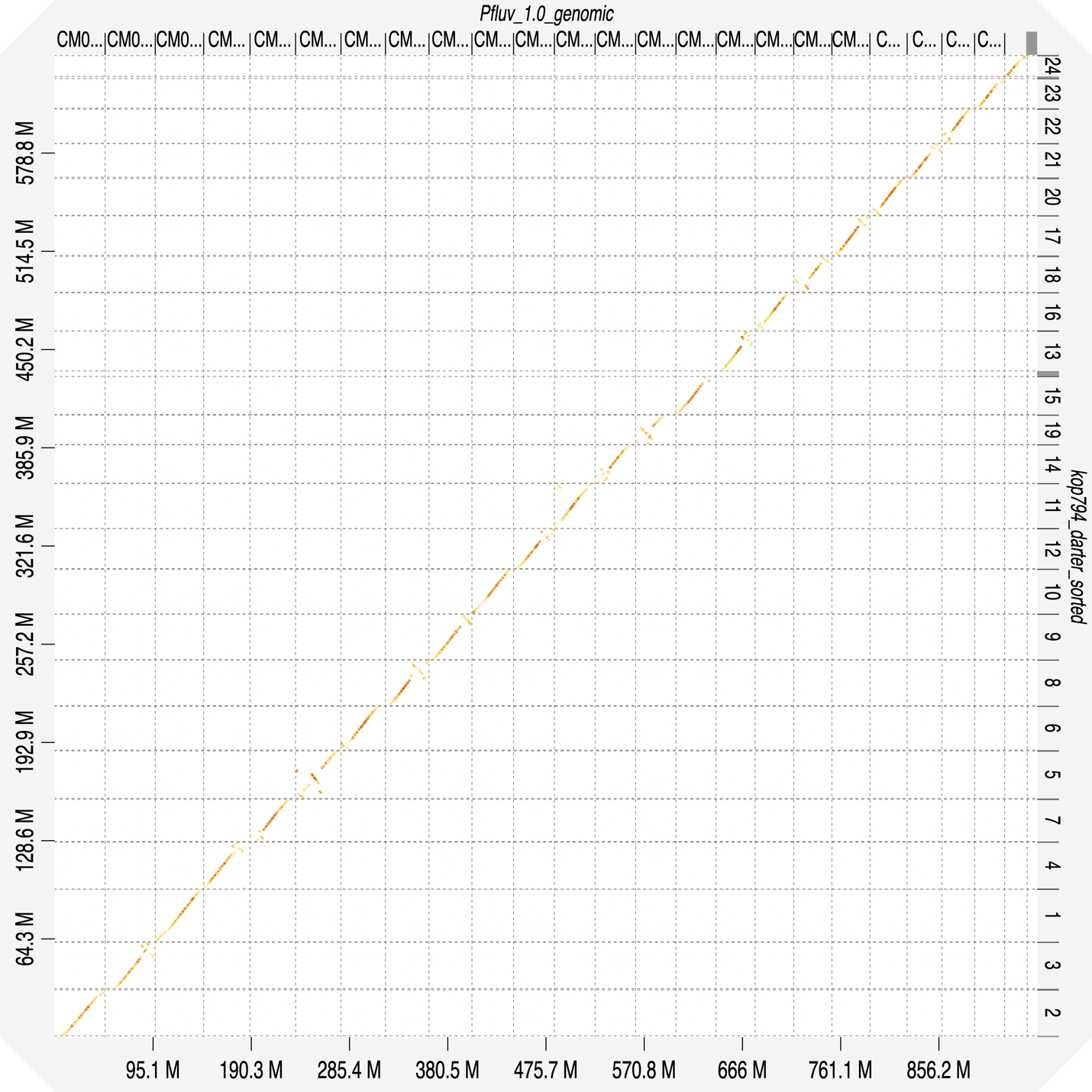

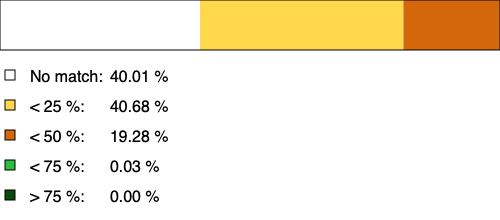

Supporting Figure 4. Proportion of total reads removed as clones by Rapture batch. Four library preparations (96-well plates) made using different adaptors were pooled in each batch, and clones were identified and removed on the plate level.

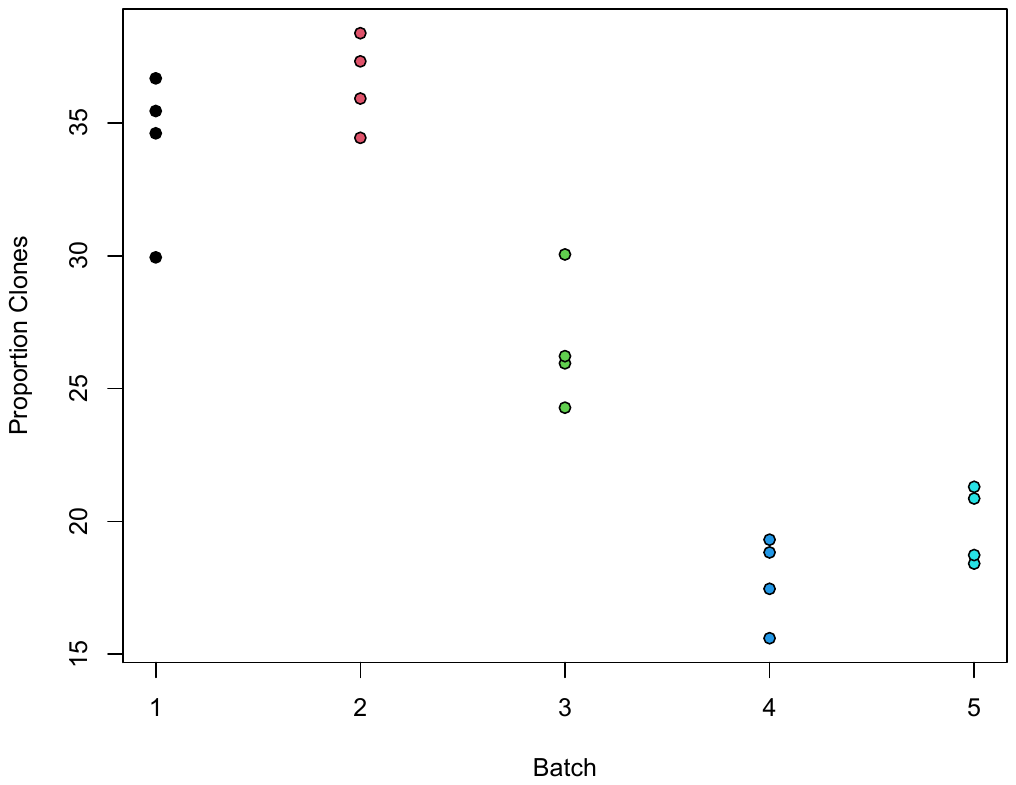

Supplemental Figure 5a. Proportion of reads mapping to the *E. cragini* reference and proportion of these mapped reads aligning to the Rapture loci for each batch and species.
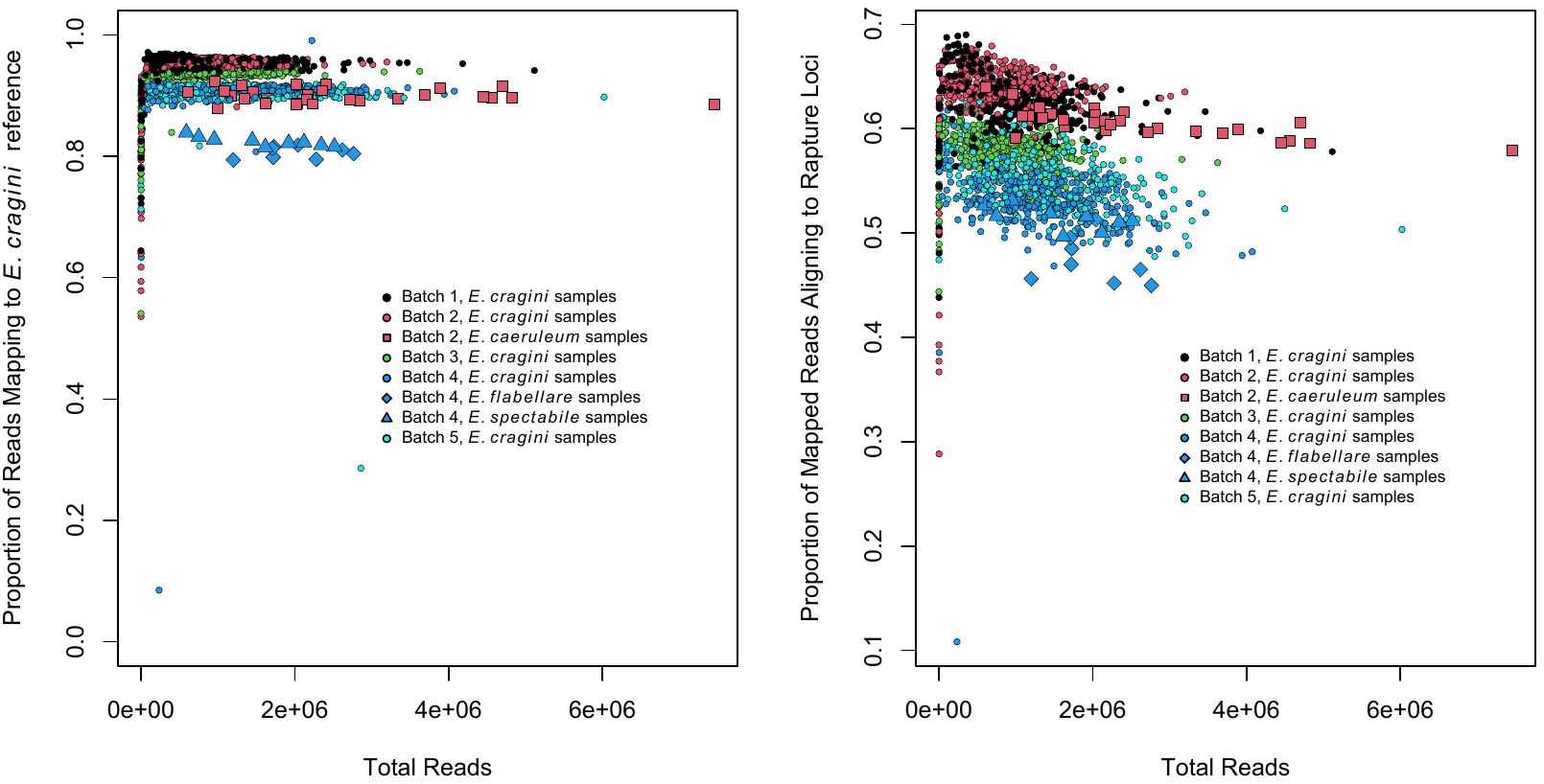

Supporting Figure 5b. Proportion of reads mapping to the *E. spectabile* reference and proportion of these mapped reads aligning to the Rapture loci for each batch and species.

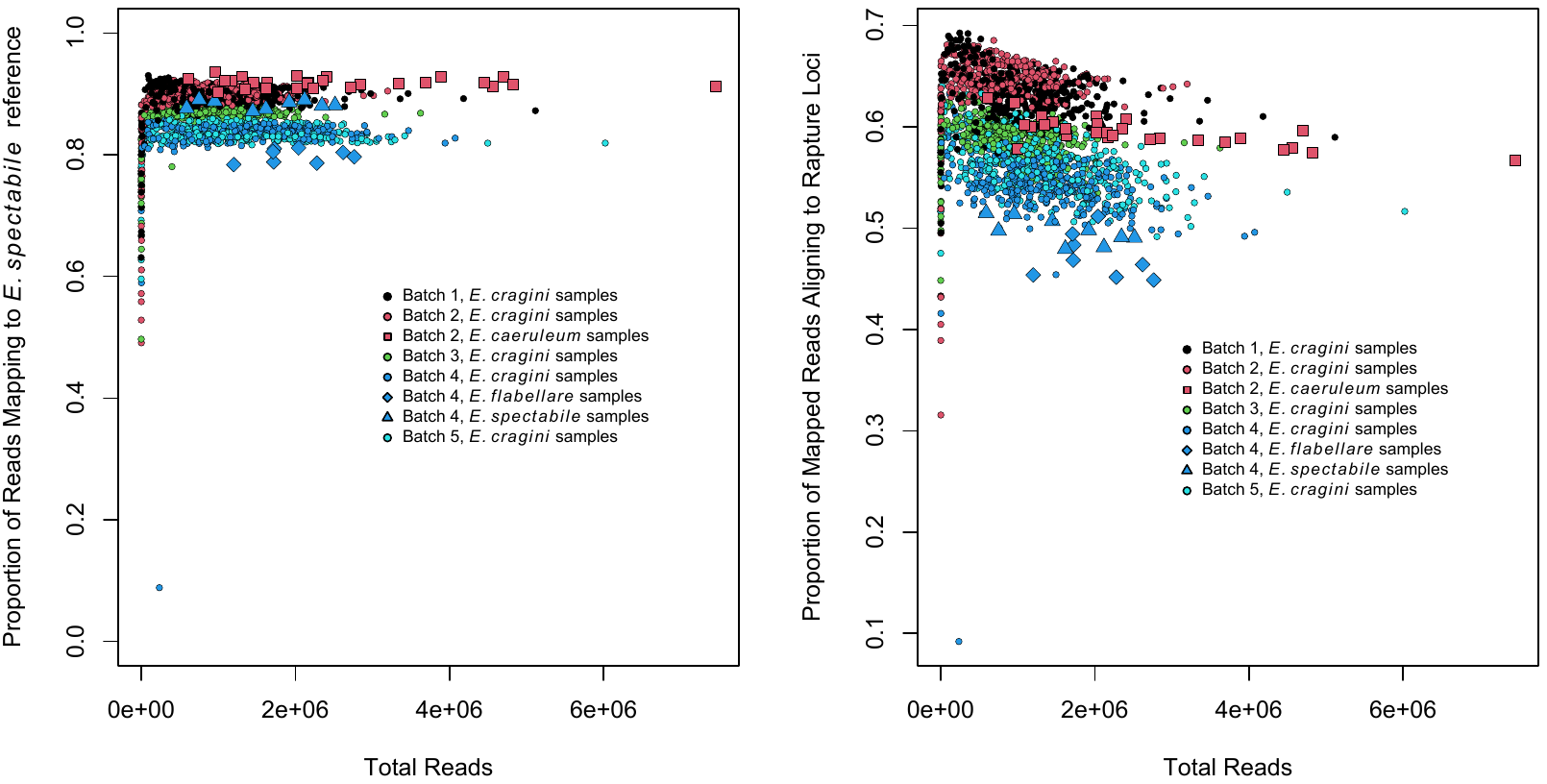

Supplemental Figure 6. Expanded coverage statistics at multiple read depths for different categories of Rapture loci.

(a) Short loci aligned to *E. cragini* reference genome.

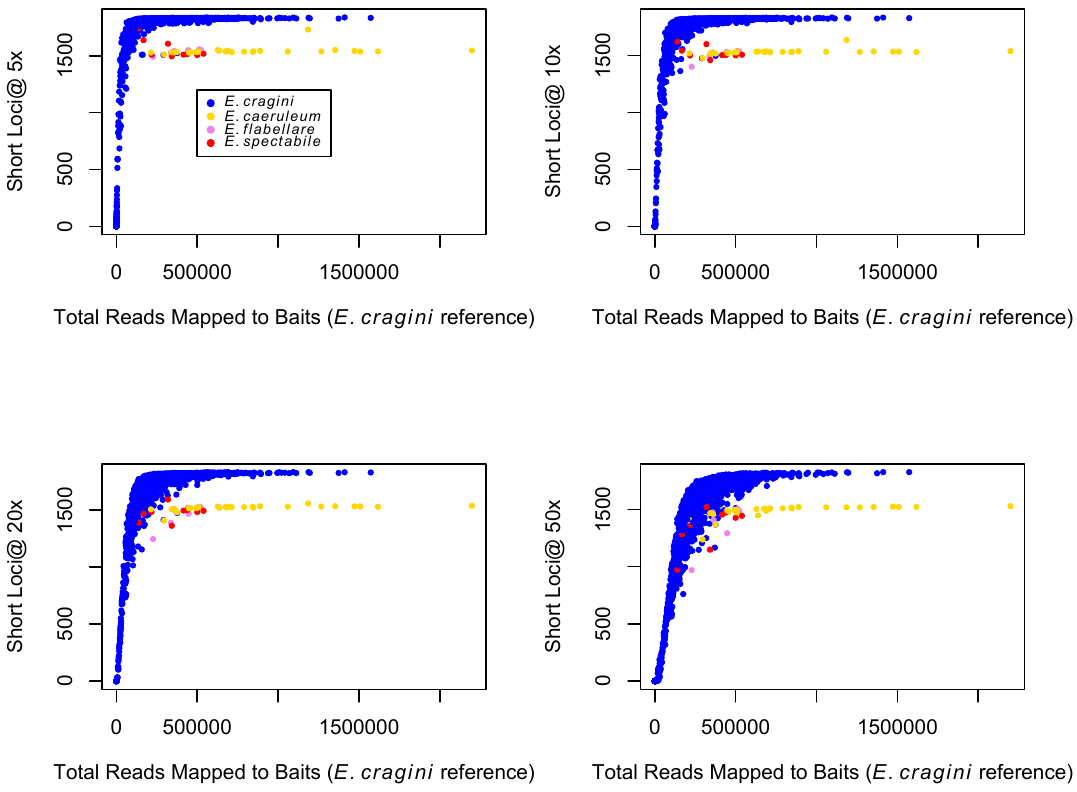

(b) Short loci aligned to *E. spectabile* reference genome.

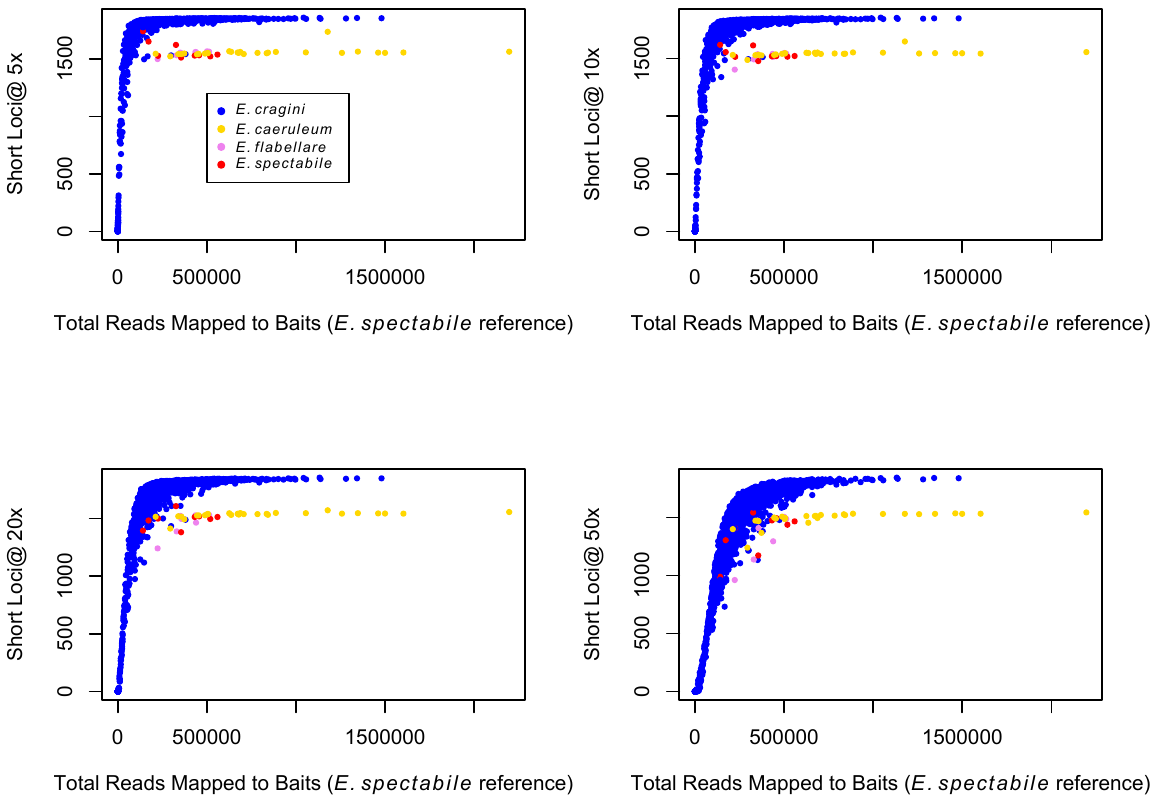

(c) Long loci aligned to *E. cragini* reference genome.

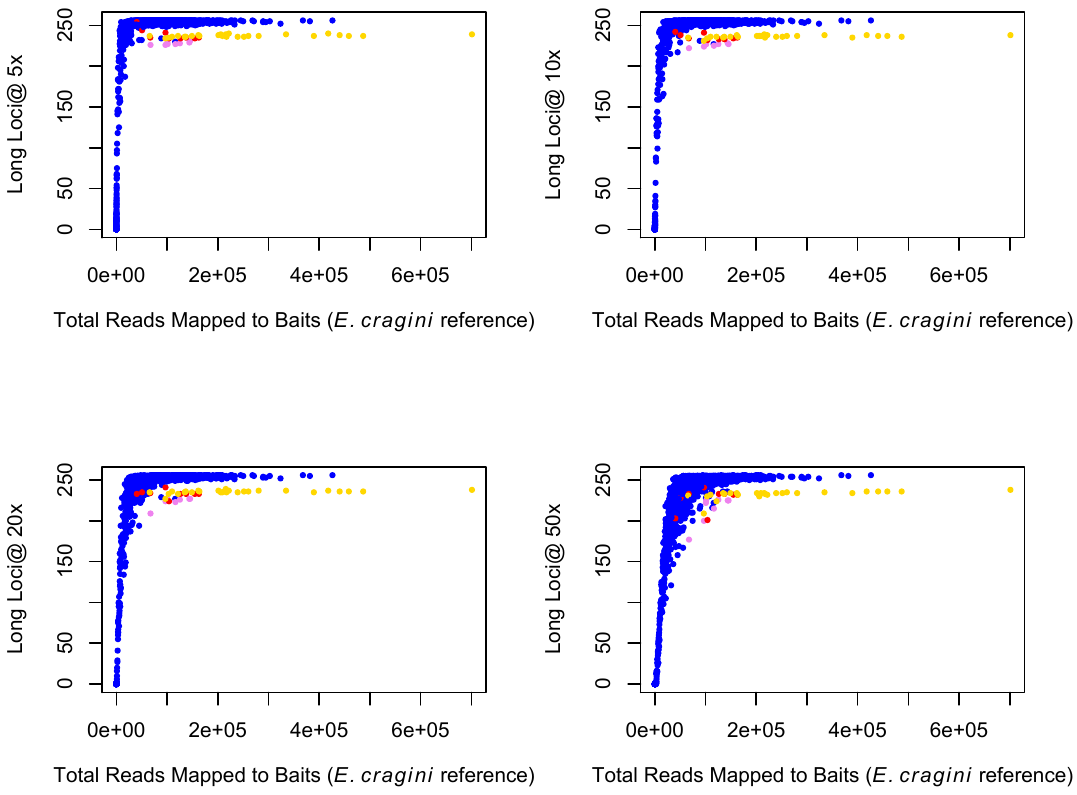

(d) Long loci aligned to *E. spectabile* reference genome.

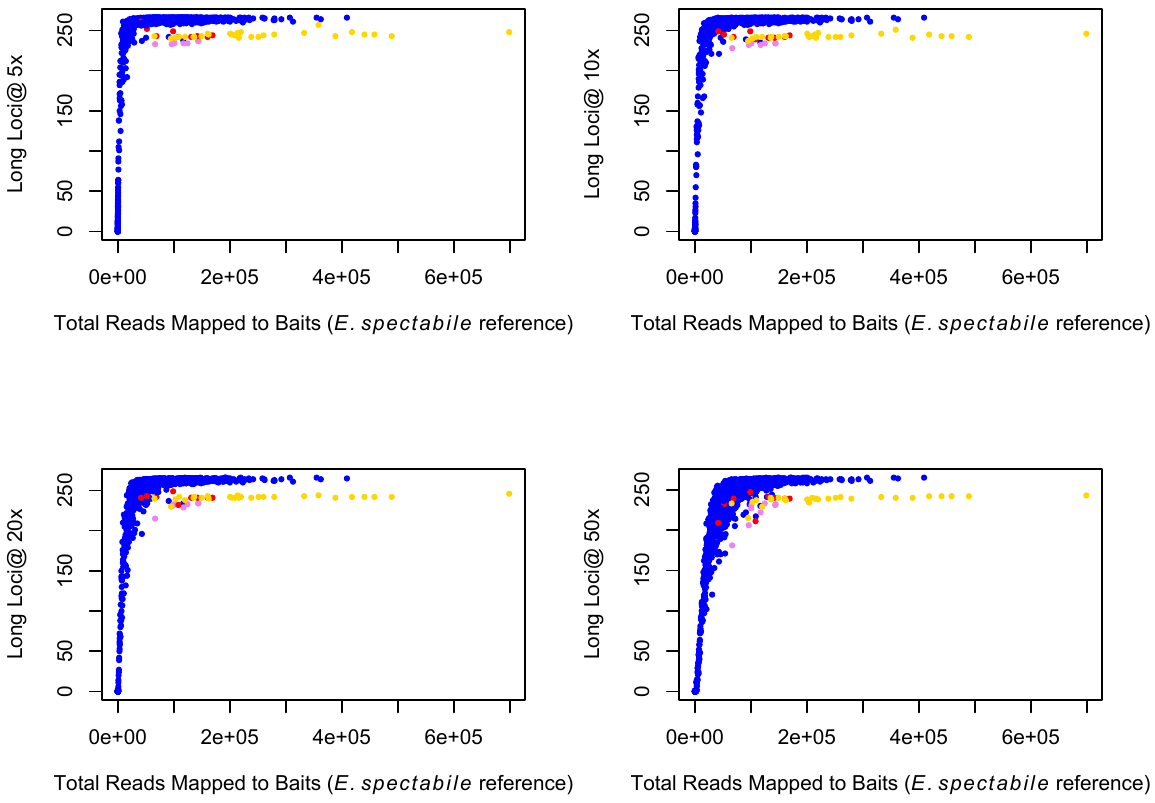

(e) Loci putatively under selection aligned to *E. cragini* reference genome.

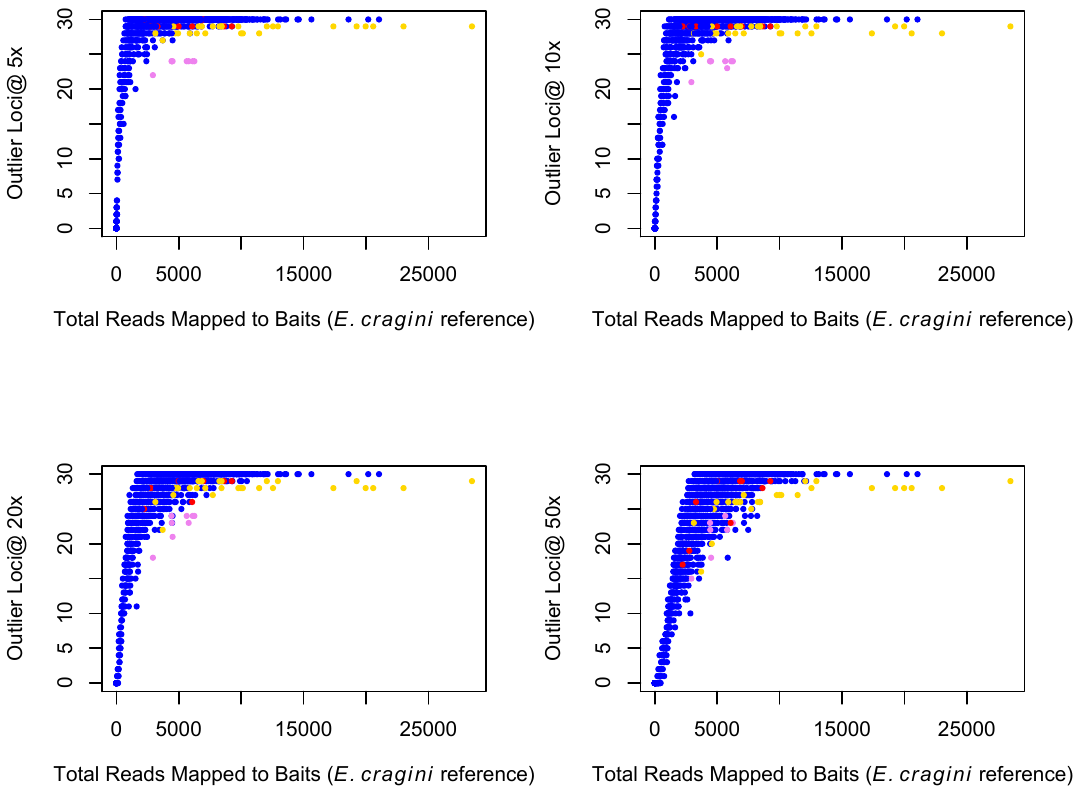

(f) Loci putatively under selection aligned to *E. spectabile* reference genome.

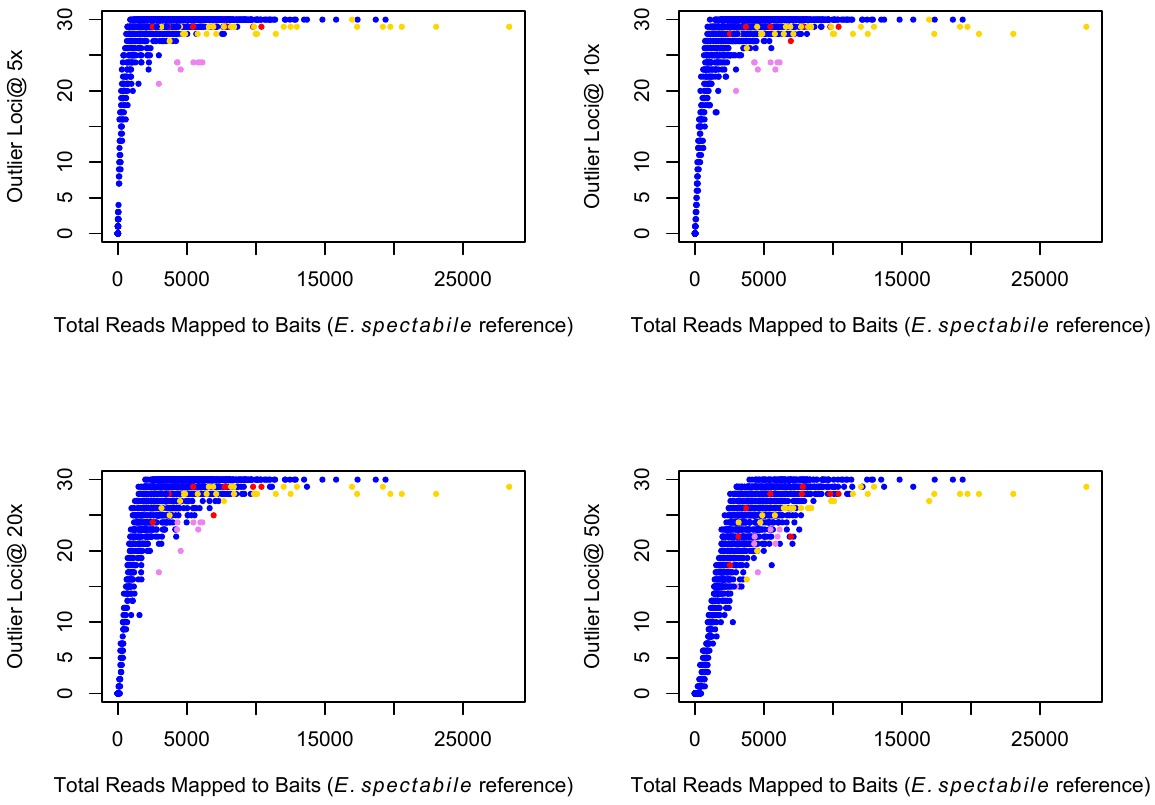

Supplemental Figure 7. Coverage for either long (top) or short (bottom) Rapture loci for a single *E. cragini* individual. Overlapping blue lines on the left panels indicate per-base coverage for all bases within the Rapture locus and a 500-bp buffer around the locus. The right panels show the number of loci for which each base in this region is covered at >20x. The locations of the restriction cut site and the baits are shown in the right panels as well. Read depth beyond the portions of the sequence covered by the capture baits shows a hump-shaped distribution corresponding to reverse reads with varying degrees of overlap created by the random shearing step. Read depth tends to drop off rapidly for short loci, although some bases up to 500 bp from the restriction site were represented at high coverage. For the long loci, however, coverage was consistently high at all bases up to approximately 500 bp in either direction from the restriction site).

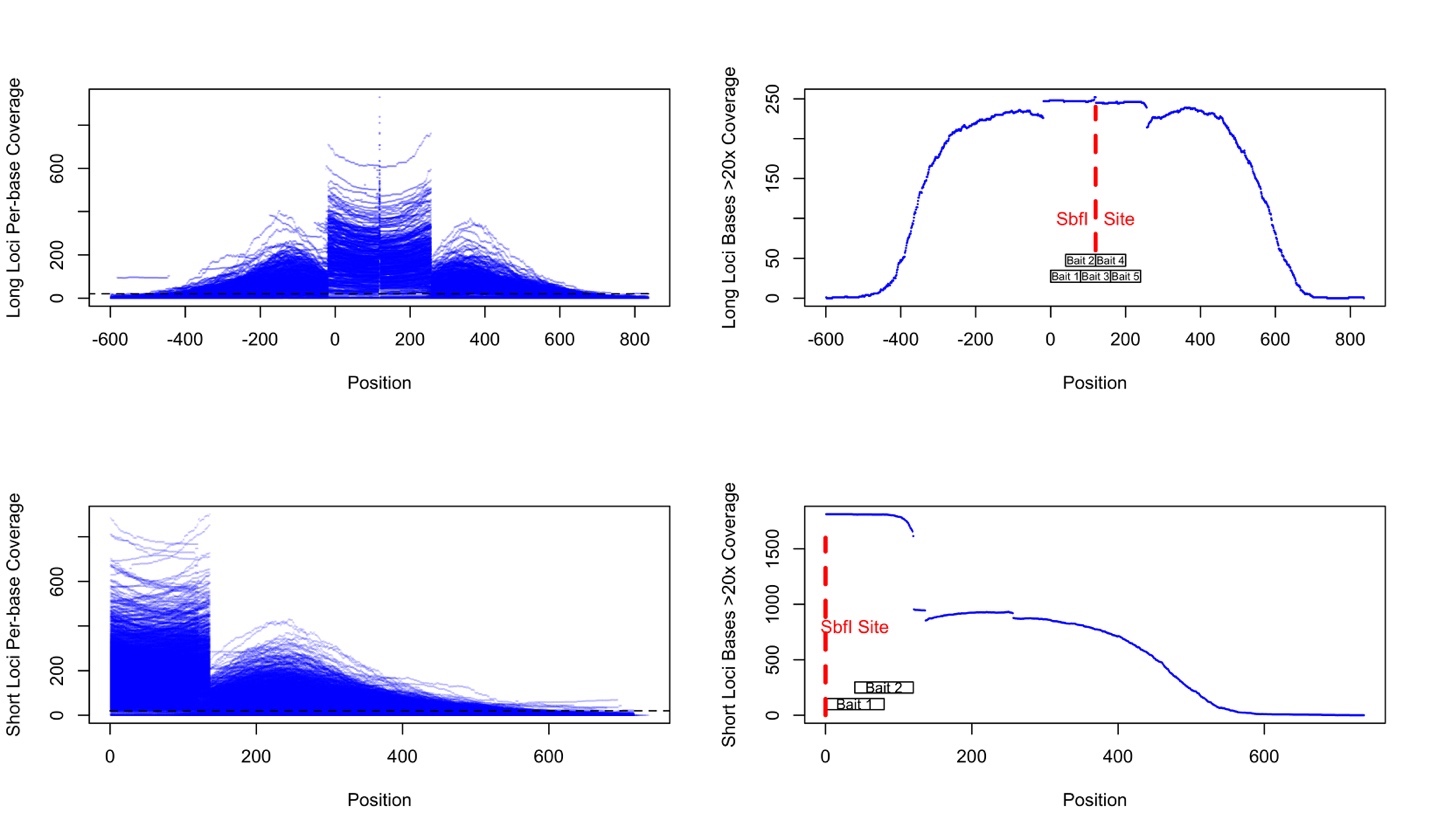

Supporting Figure 8a. PCA results (with points colored either by batch or metapopulation) for full Rapture dataset aligned to *E. cragini*.

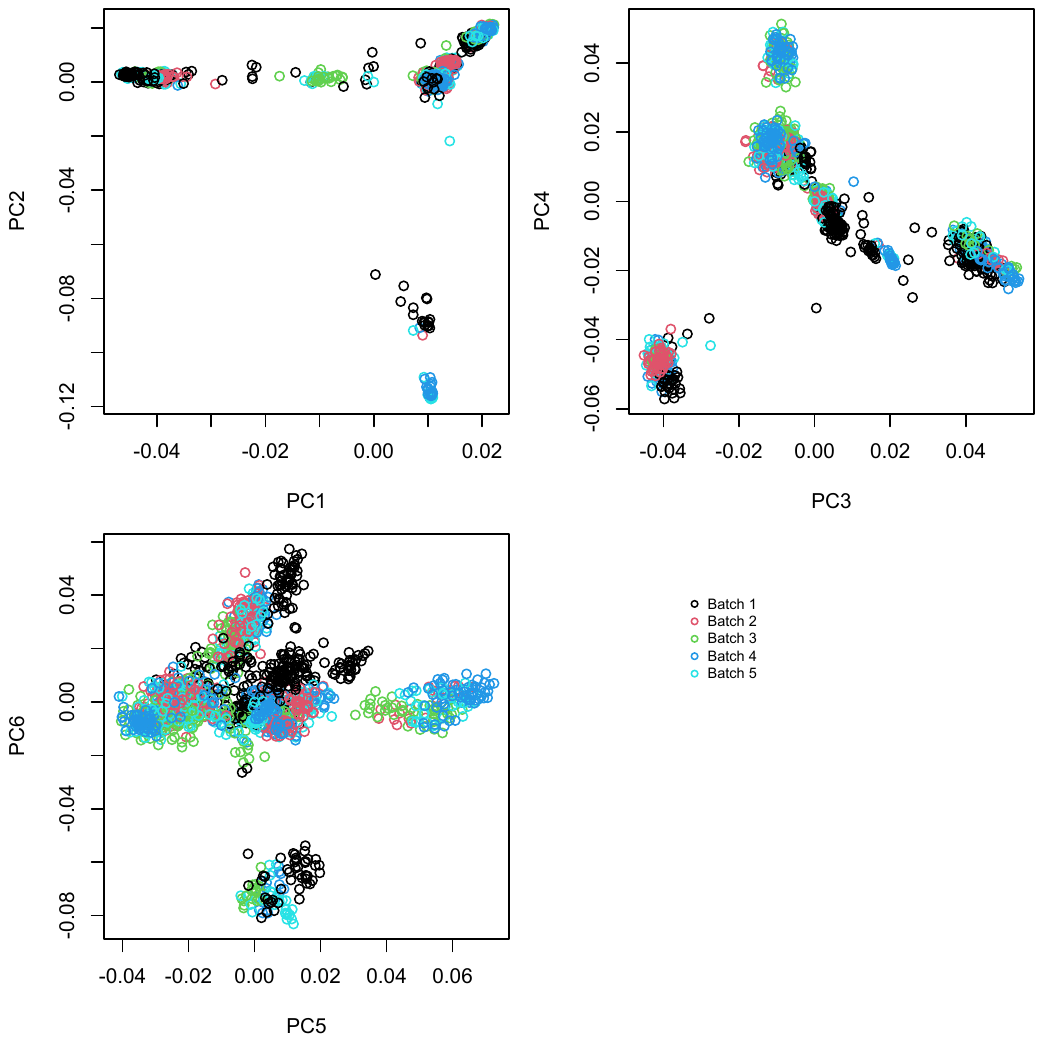

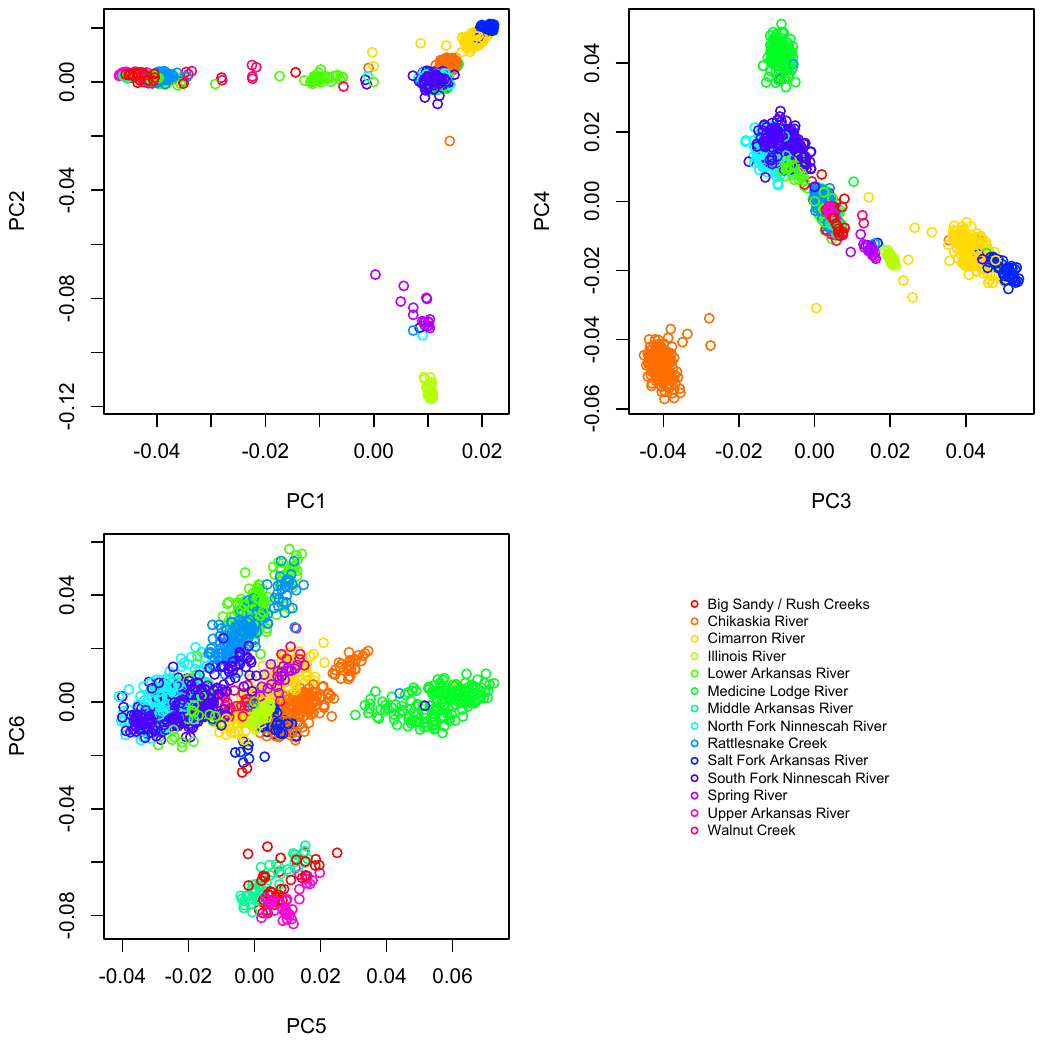

Supporting Figure 8b. PCA results (with points colored either by batch or metapopulation) for full Rapture dataset aligned to *E. spectabile*.

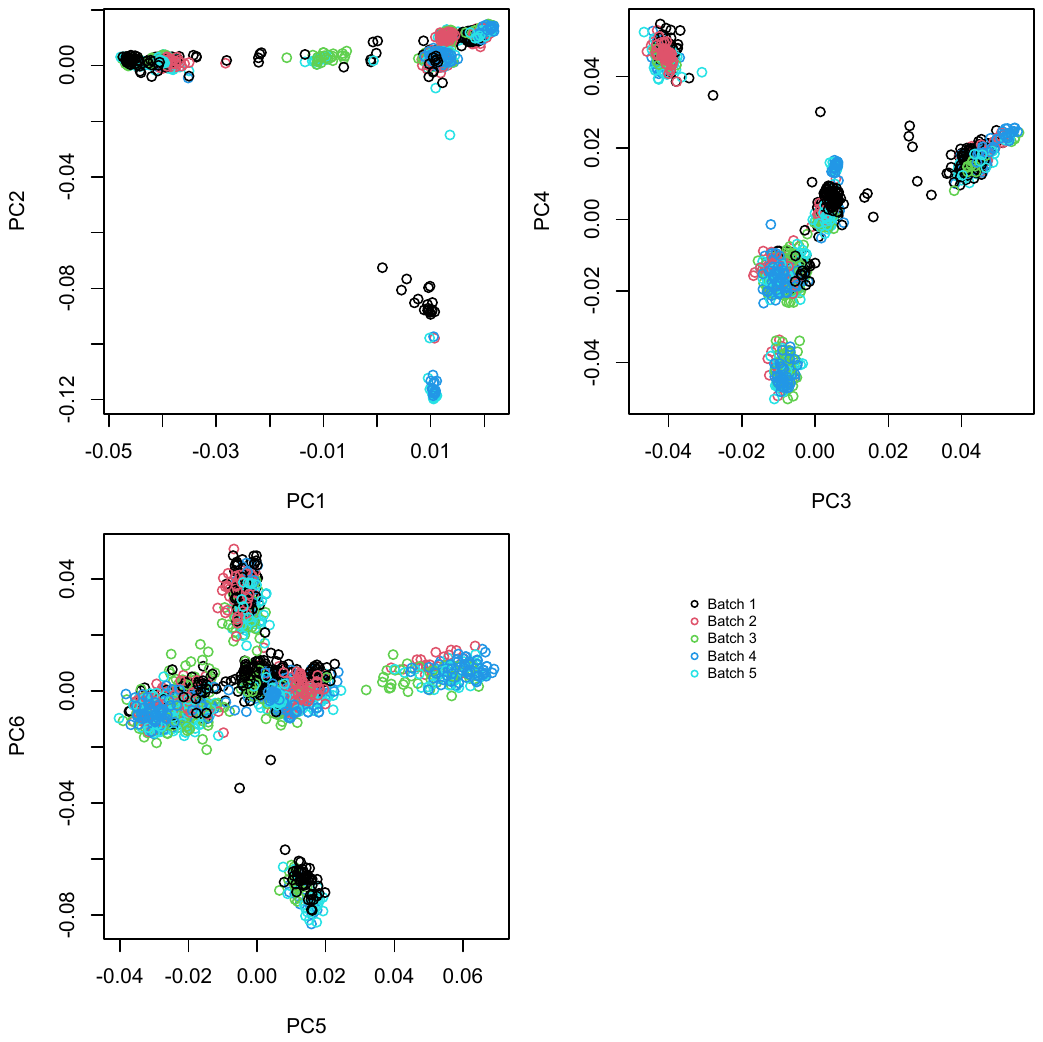

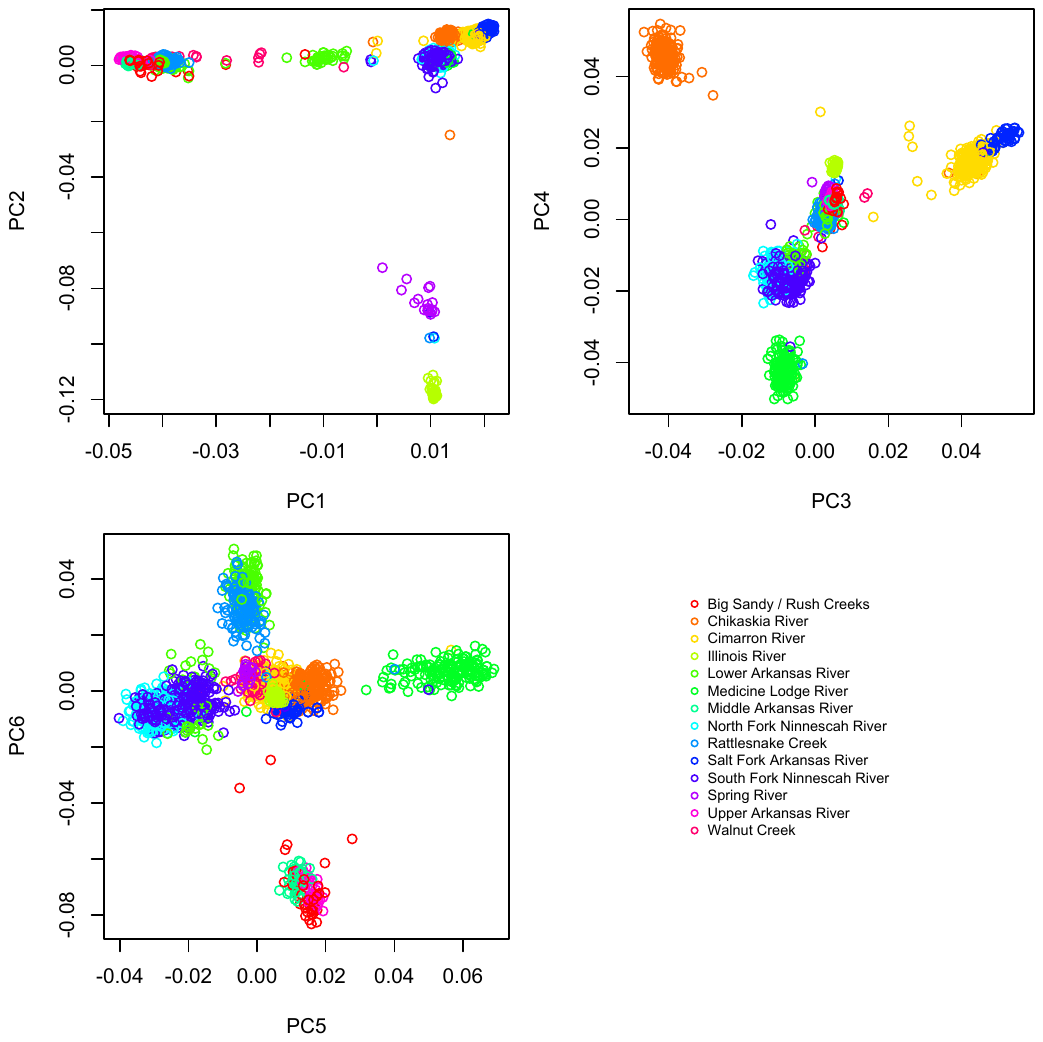

Supporting Figure 8c. PCA results (with points colored either by batch or metapopulation) for subsampled Rapture dataset aligned to *E. cragini*.

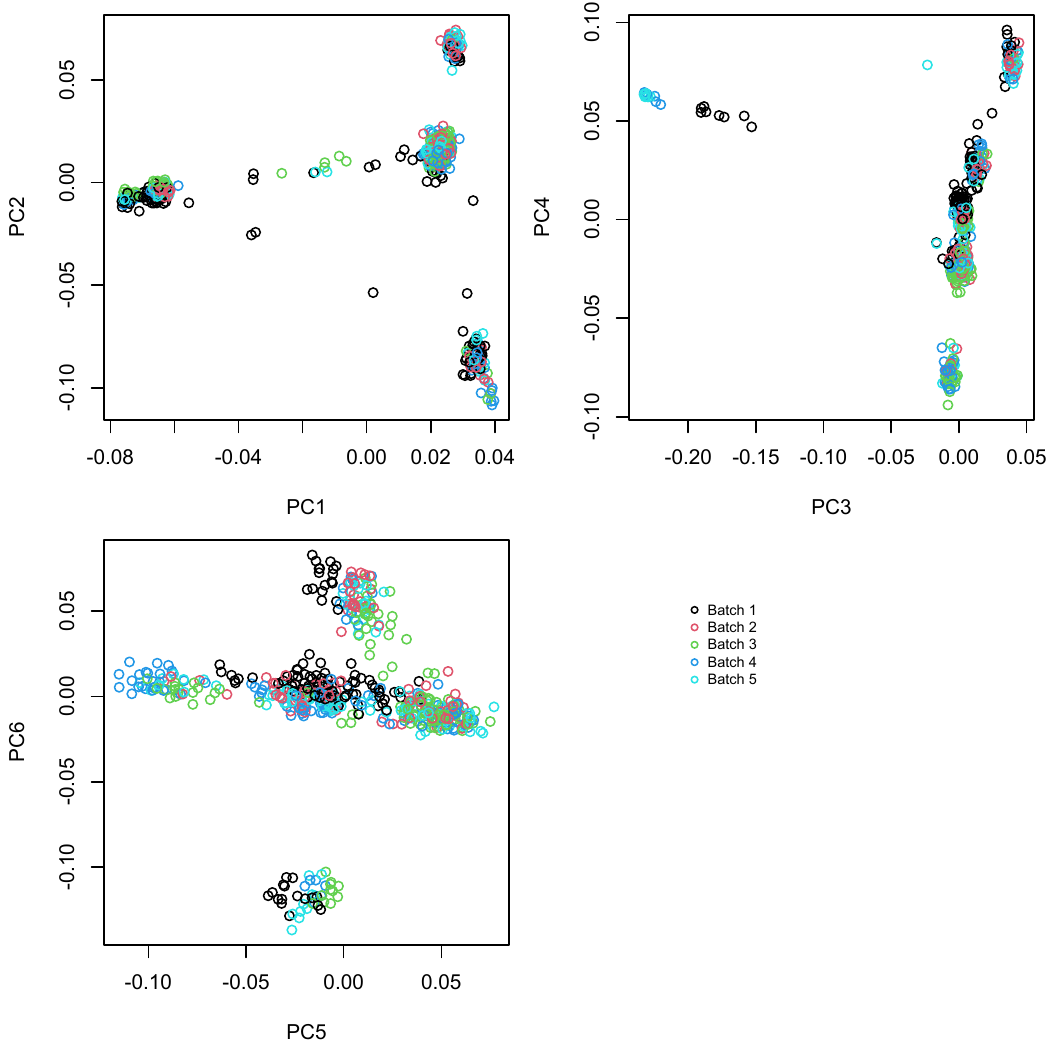

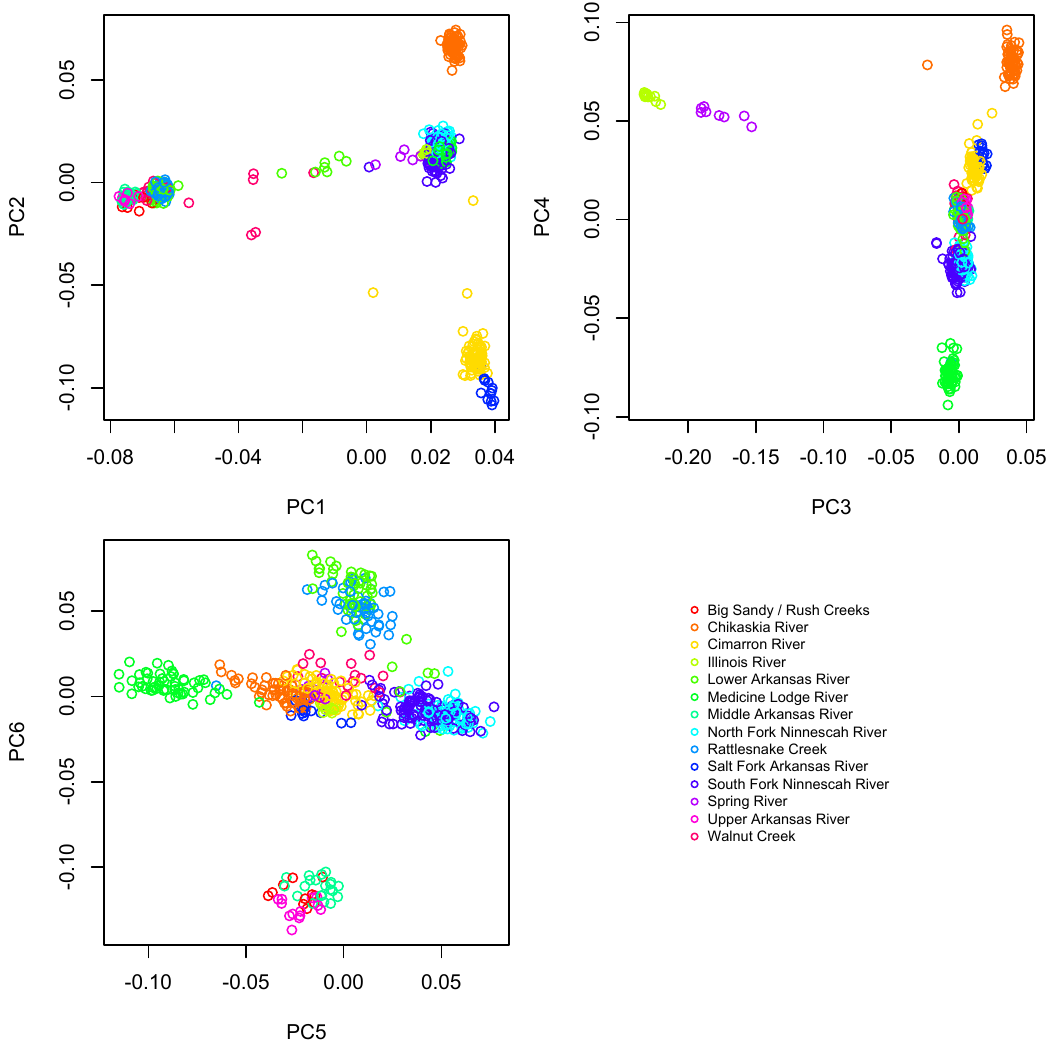

Supporting Figure 8d. PCA results (with points colored either by batch or metapopulation) for subsampled Rapture dataset aligned to *E. spectabile*.

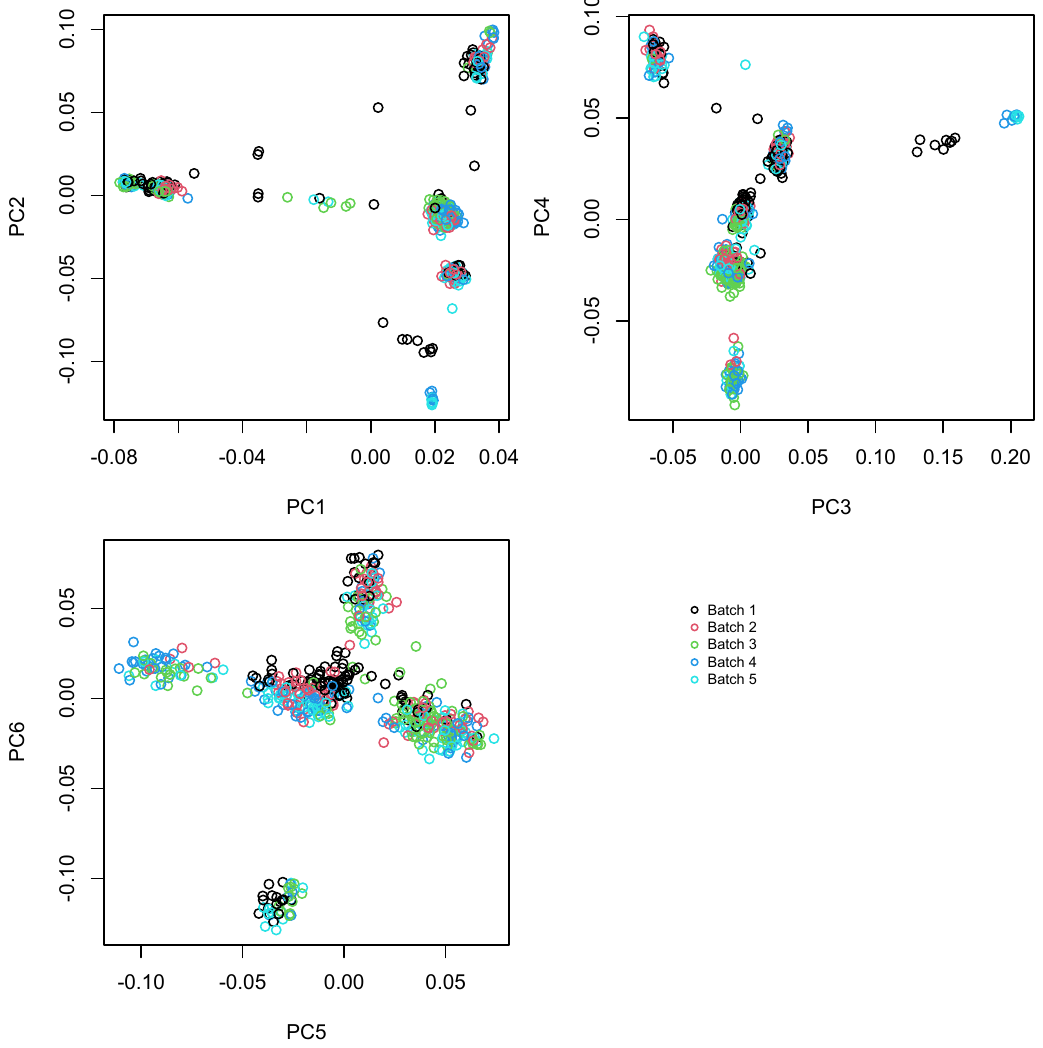

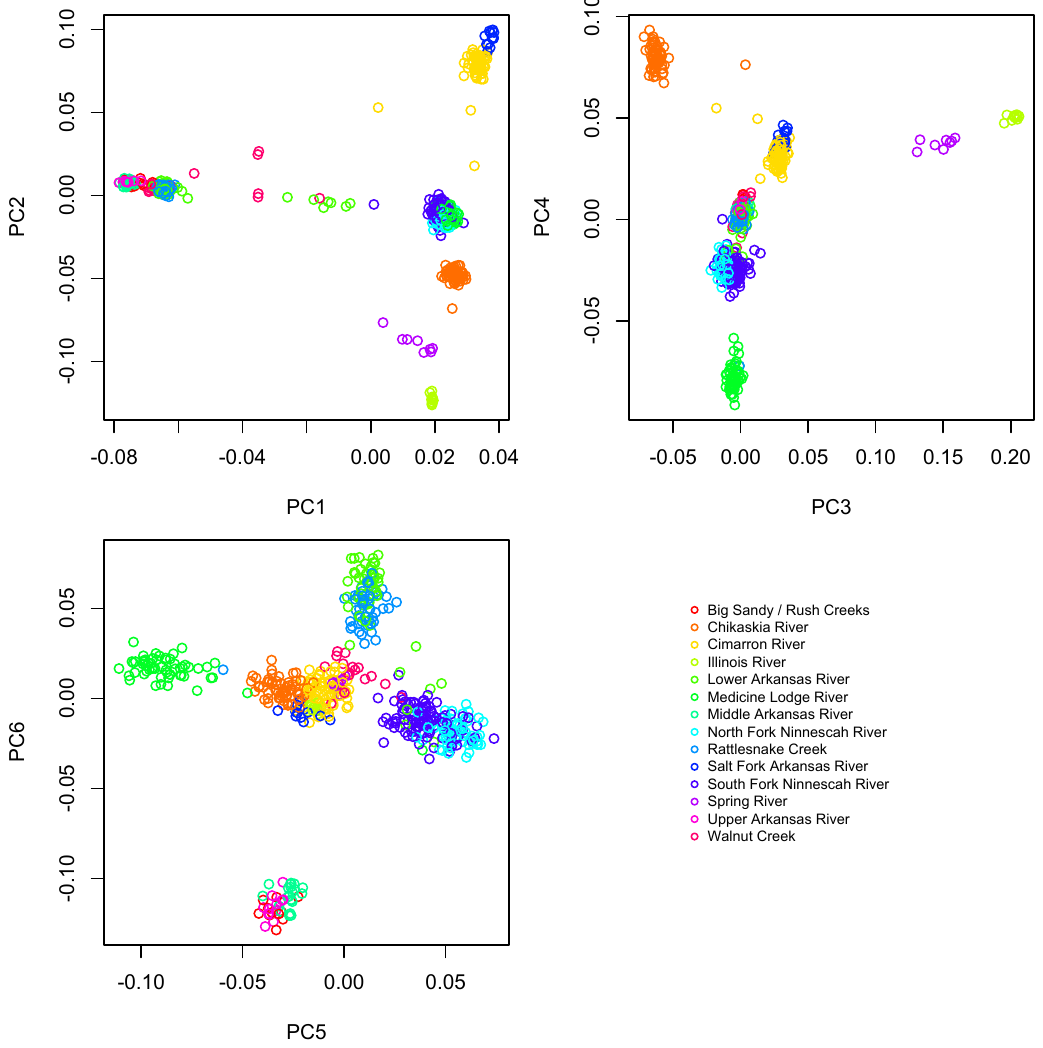

Supporting Figure 9. Additional population structure figures.

(a) Rapture loci, subsetted dataset, *E. cragini* reference
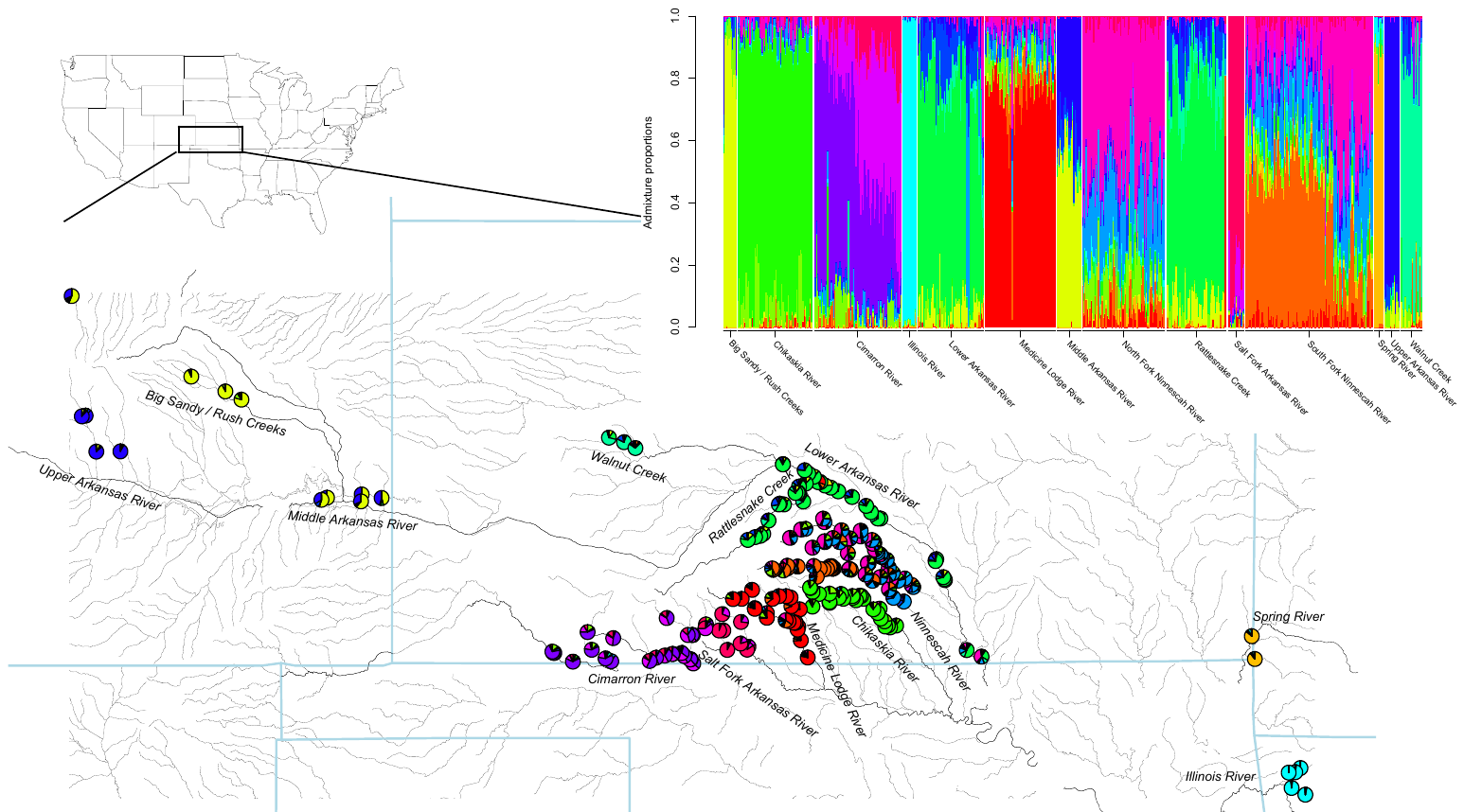

(b) Rapture loci, subsetted dataset, *E. spectabile* reference
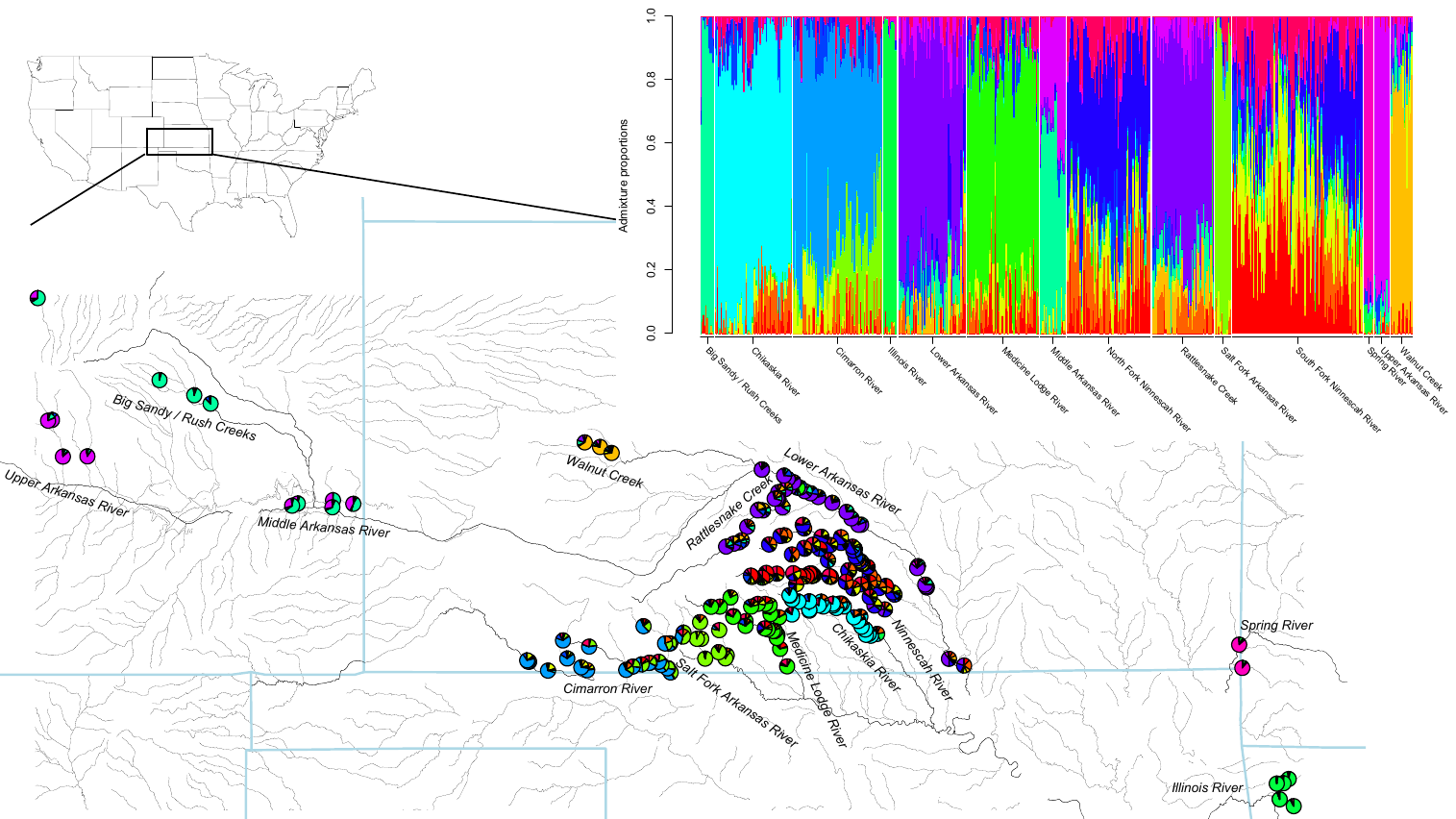

(c) Rapture loci, full dataset, *E. spectabile* reference

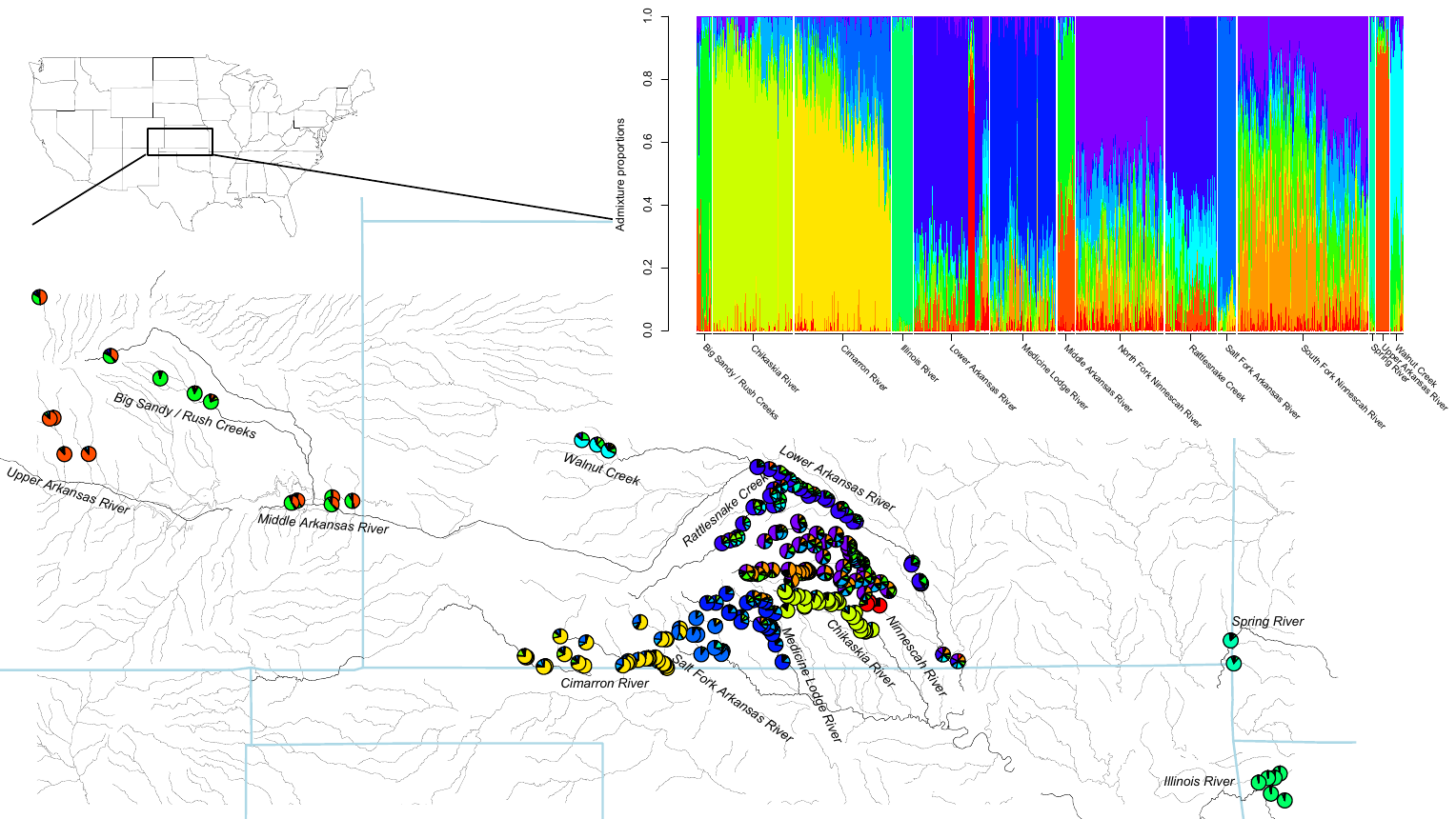

(d) WGS dataset, *E. spectabile* reference

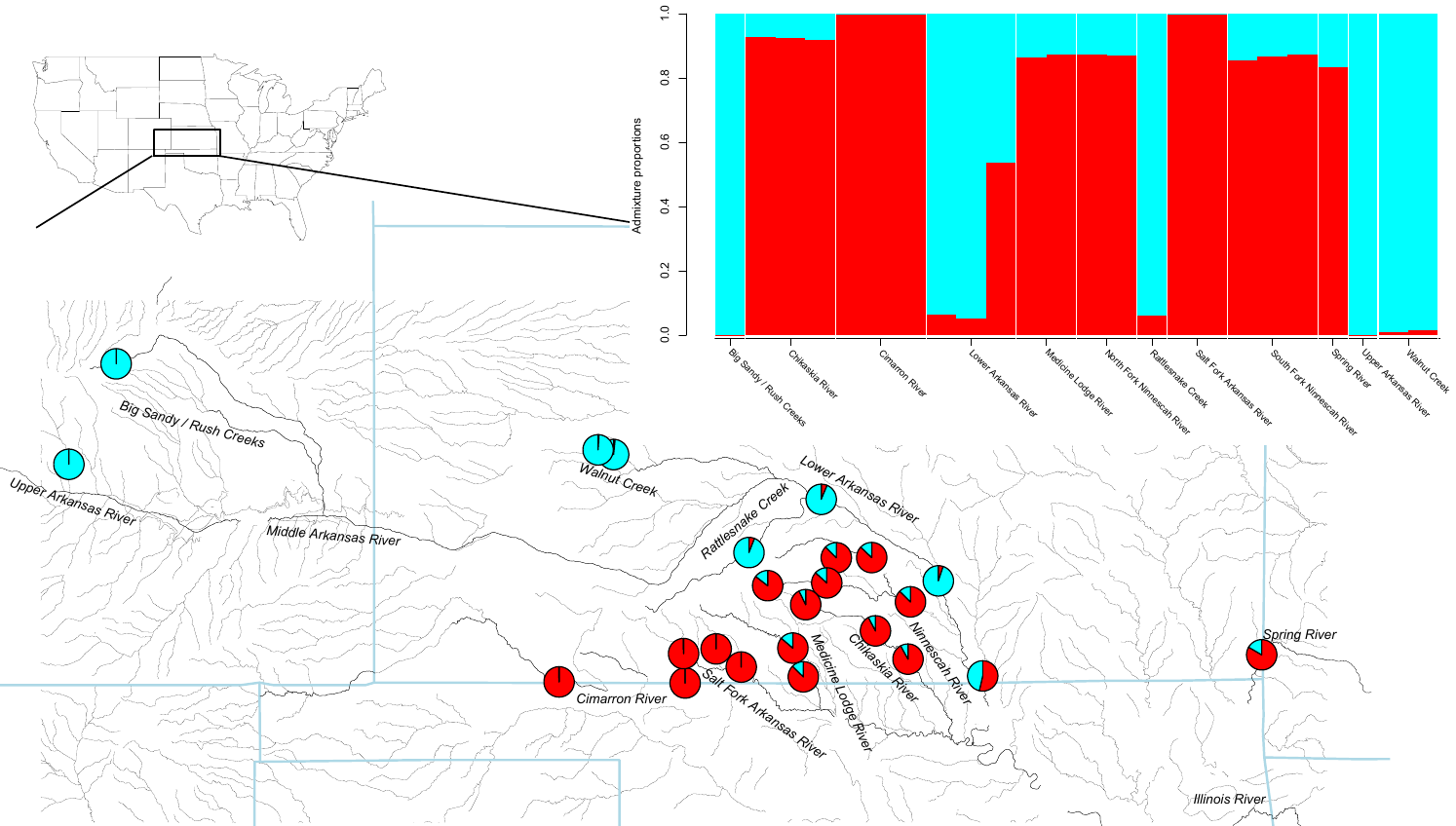

Supporting Figure 10. Selection scan statistics calculated in PCAngsd using the first principal component for Rapture and WGS datasets.

(a) Datasets aligned to the *E. cragini* reference genome.

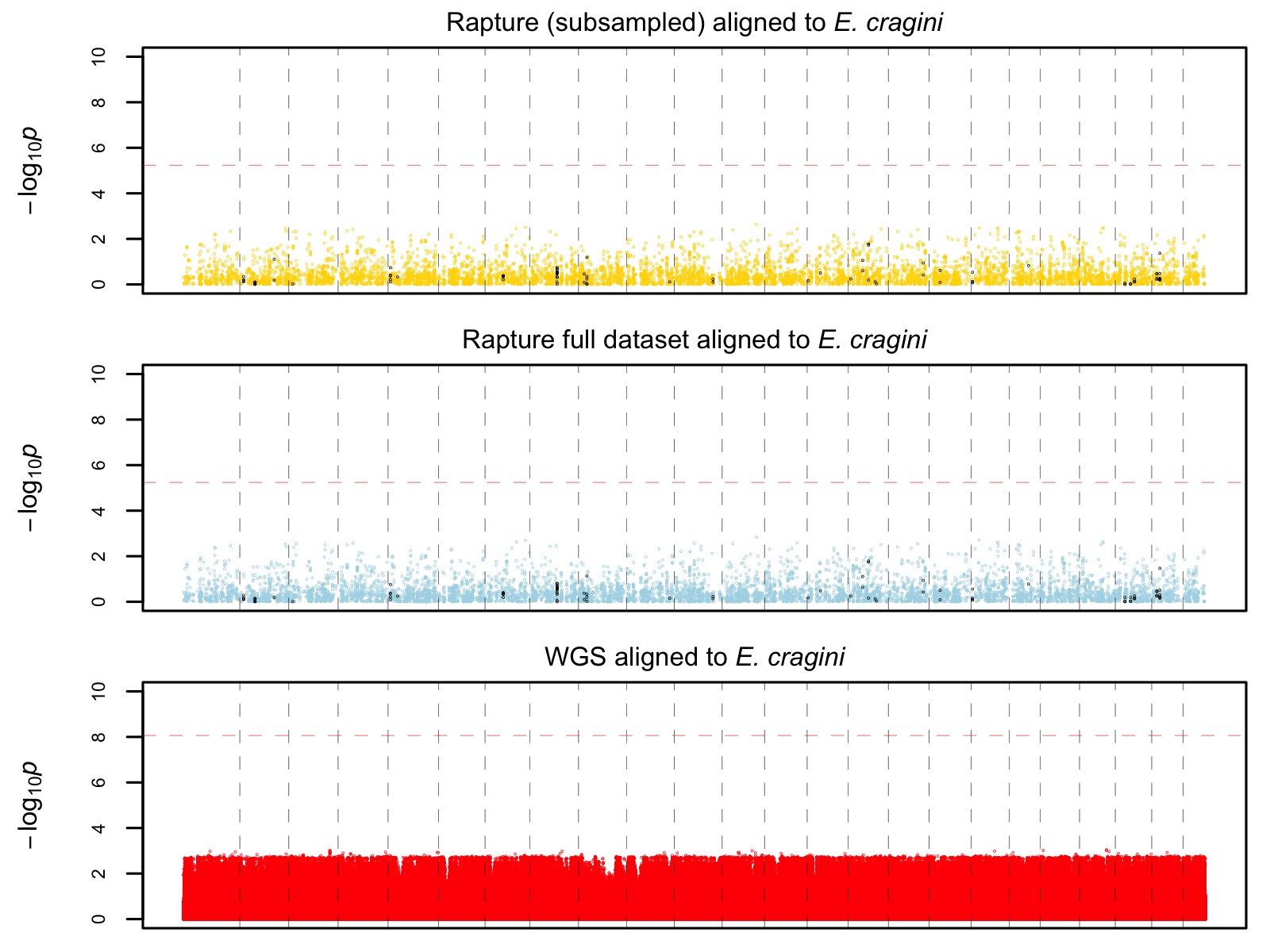

(b) Datasets aligned to the *E. spectabile* reference genome.
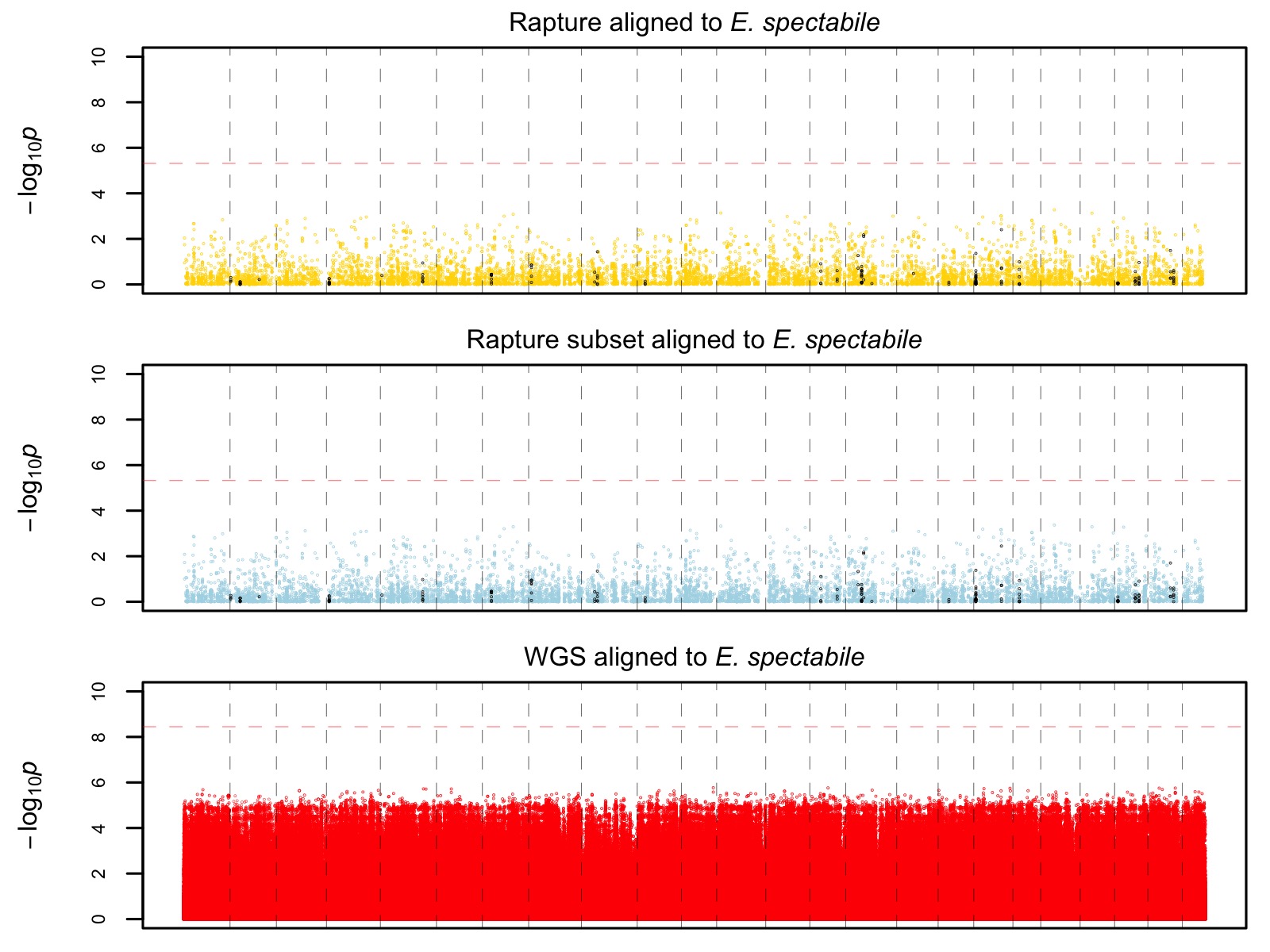

Supporting Figure 11. Phylogenetic informativeness profiles for short (NS), long (NL), and selected (SB) baits.
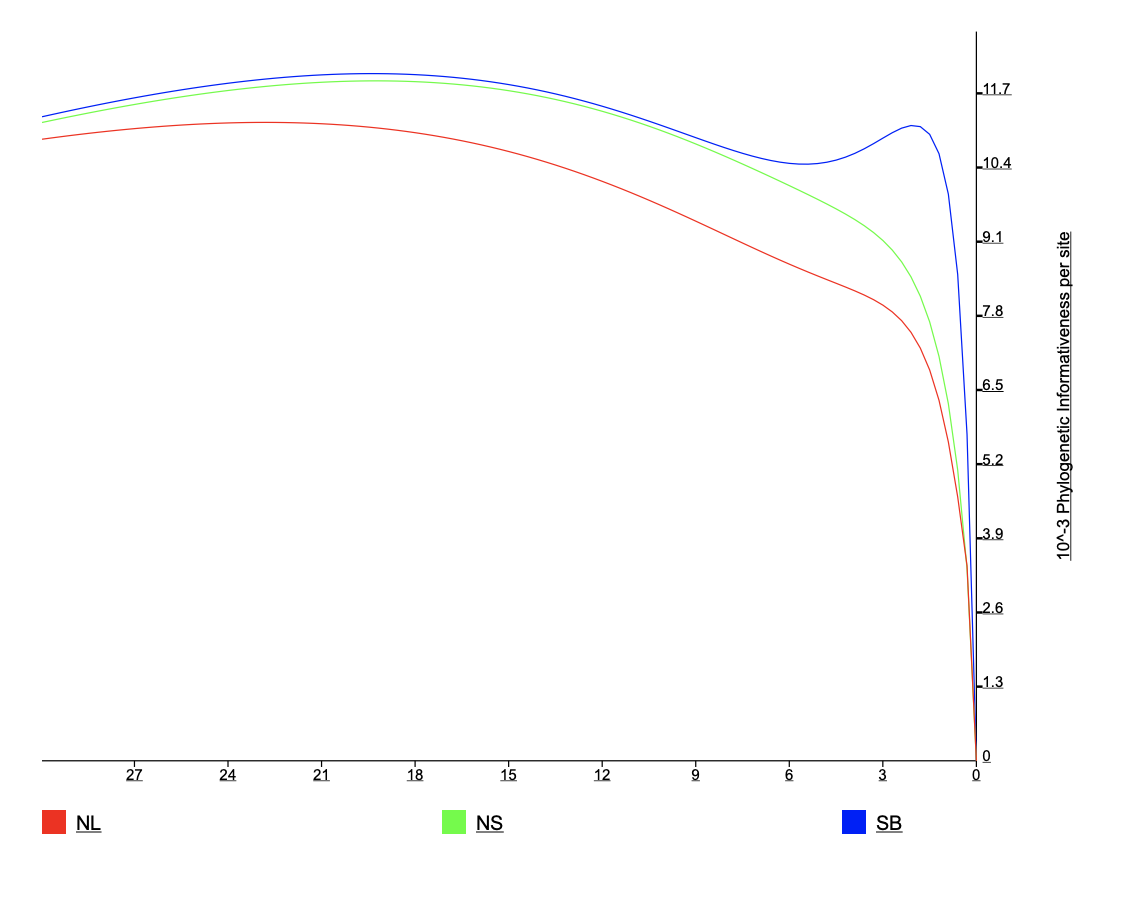
